# Mammalian mitochondrial polyphosphate is regulated by the 5-InsP_7_ synthase IP6K1

**DOI:** 10.1101/2025.06.17.659843

**Authors:** Jayashree S. Ladke, Azmi Khan, Anshit Singh, Jyoti Singh, Nicole Steck, Henning J. Jessen, Ullas Kolthur-Seetharam, Manish Jaiswal, Rashna Bhandari

## Abstract

Inorganic polyphosphate (polyP) is a linear polymer of varying length composed of phosphate residues linked by phosphoanhydride bonds. PolyP remains poorly understood in mammals due to its low abundance and lack of information on its synthesis and regulation. Using synthetic polyP_8_ as a bait, we identified polyP interacting proteins involved in RNA splicing, transcription, and mitochondrial function. We developed a DAPI fluorescence-based assay to quantify the low levels of polyP present in mammalian cell lines and tissues, and detected a higher concentration of polyP in the mitochondria compared with the nucleus and post-mitochondrial fraction. We observed that mitochondrial polyP synthesis relies on active FoF1 ATP synthase and an intact proton gradient across the inner mitochondrial membrane. Our data shows that orthophosphate (Pi) is required for the generation of mitochondrial polyP and that the presence of ATP enhances Pi-driven polyP synthesis in isolated mitochondria. We discovered that the inositol pyrophosphate 5-InsP_7_, synthesized by IP6K1, regulates mitochondrial polyP levels. Mice and cells deficient in IP6K1 showed a significant reduction in mitochondrial polyP synthesis compared with wild type controls. Cells lacking IP6K1 also showed impaired mitochondrial respiration and membrane potential. The expression of active IP6K1, but not its catalytically inactive form, restored mitochondrial polyP synthesis and membrane potential in IP6K1 deficient cells, but mitochondrial respiration was rescued by expression of either active or inactive IP6K1. These data show that IP6K1 regulates mitochondrial function and polyP production through both the synthesis of 5-InsP_7_ and via a catalytic activity-independent mechanism. Our findings uncover a link between 5-InsP_7_, an energy sensor, and polyP, an energy store, in the regulation of mammalian mitochondrial homeostasis.

## INTRODUCTION

Inorganic polyphosphate (polyP) is a linear polymer of orthophosphate residues linked by high-energy phosphoanhydride bonds. This prebiotic polymer is ubiquitous across all life forms, varying in its chain length, localisation, concentration and function across different model organisms studied till date [1–3]. In mammals, polyP has been detected in different tissues like brain, heart, kidney liver, and lung [4], cell types like fibroblasts [5], myeloma cells [6], and platelets [7], and subcellular compartments like nucleus, mitochondria, plasma membrane, granules, and lysosome-related organelles [4, 5]. Mammalian polyP, while low in abundance compared with unicellular eukaryotes, participates in several biological processes including blood clotting [7, 8], mitochondrial calcium buffering [9, 10], bone mineralization [11] and energy metabolism [12].

The synthesis of polyP and its regulation have been studied extensively in bacteria and unicellular eukaryotes. In bacteria, polyP is synthesized by polyphosphate kinase 1 (PPK1) [13], whereas in budding yeast, polyP is synthesized by the vacuolar transport chaperone (VTC) complex [14], a multi-subunit, membrane-bound complex that synthesizes and sequesters polyP into the lumen of vacuoles. To date, no definitive polyP synthase has been identified in mammals. Previous work has shown that isolated mitochondria from rat liver show polyP synthesis on incubation with substrates of the electron transport chain (ETC) [15–17], and it was suggested that FoF1 ATP synthase may be a polyP synthase in mammalian mitochondria.

In yeast, the catalytic activity of the VTC complex is allosterically activated by the inositol pyrophosphate 5-diphosphoinositol pentakisphosphate (5-InsP_7_), a phosphate-rich signalling molecule derived from *myo*-inositol substituted by monophosphate and diphosphate moieties [18–20]. The *S. cerevisiae* mutant *kcs1*Δ, which lacks the IP6 kinase that synthesizes 5-InsP_7_ from InsP_6_, has undetectable levels of polyP. In mammals, 5-InsP_7_ is synthesised by inositol hexakisphosphate kinase (IP6K), which has 3 paralogs -IP6K1/2/3. We have previously reported a substantial decrease in polyP levels in the platelets of *Ip6k1^-/-^* mice compared with *Ip6k^+/+^* mice [21]. Reduced polyP release during platelet degranulation in *Ip6k1^-/-^* mice was shown to result in impaired blood clotting [21], and reduced formation of neutrophil-platelet aggregates, alleviating inflammation-associated lung damage during a bacterial challenge [22]. These studies demonstrated that the relationship between cellular levels of InsP_7_ and polyP is maintained in yeast and mammals despite the lack of conservation in the mechanism of polyP synthesis in these classes of organisms.

Our present study focuses on understanding the relationship between 5-InsP_7_ and polyP in mammalian mitochondria. We show that mitochondrial polyP levels depend directly on mitochondrial activity. Inhibition of mitochondrial respiration reduces mitochondrial polyP levels as well as the ability of isolated mitochondria to synthesize polyP. Our data confirms earlier studies suggesting that orthophosphate (Pi), and not ATP, serves as the source of phosphate for mammalian mitochondrial polyP synthesis. Intriguingly, ATP hydrolysis supports Pi-dependent polyP synthesis in isolated mammalian mitochondria. We show that IP6K1 maintains mitochondrial polyP levels and mitochondrial membrane potential via its ability to catalyze the synthesis of 5-InsP_7_. Additionally, IP6K1 is required for the maintenance of mitochondrial respiration independent of its catalytic activity. Our data provides the first evidence of a direct link between mitochondrial polyP, believed to be a store for Pi in mitochondria during times of ATP sufficiency, and 5-InsP_7_, the metabolic sensor that is sensitive to cellular ATP levels.

## RESULTS

### The mammalian polyP interactome includes proteins of the mitochondria

Previous studies have used a protein array or sequence prediction methods to identify human and yeast proteins that interact with polyP [23–25]. We used an unbiased proteomics approach using synthetic polyP_8_ [26] conjugated with agarose beads as a bait to identify the polyP interactome in HEK293T cells, a commonly used immortalised human cell line (Fig. 1A). As negative controls, we used polyP_3_ or Pi conjugated beads, or hydroxy beads alone. The pull-down and mass spectrometry based proteome identification was conducted in replicate sets for each sample type (Table S1). By combining both replicates, we detected 453 proteins that interact with polyP_8_ but not with polyP_3_ or other control beads (Fig. 1B). Of these 109 proteins were present in both replicates of polyP_8_ (Fig. 1C). This data was subjected to the CRAPome analysis pipeline (https://reprint-apms.org/) to distinguish between authentic and contaminant interactors by comparing the total peptide numbers for each interacting protein bound to polyP_8_-agarose with the control beads (Table S2). Previously identified polyP binding proteins were present in our list of polyP_8_ interacting proteins in HEK293T cells (Table S2). We set the stringent fold change FC-B score [27] to ≥1.5, and subjected 423 proteins that fall in this range to Gene Ontology (GO) term enrichment analysis using the DAVID tool (https://david.ncifcrf.gov) [28]. PolyP binding proteins were enriched in several biological processes, notably RNA splicing and transcription by RNA polymerase II (Fig. 1D, Table S3). Among cellular component GO terms, polyP interacting proteins were present in the spliceosome, ribosome, autophagosome, and mitochondria. To validate our use of Poly_8_ beads to identify the polyP interactome, we used the heat shock protein HSP90α/β, which has previously been shown to interact with polyP [24]. Biotinylated mixed chain length polyP of ∼100 units (polyP_100_) immobilized on streptavidin agarose beads and incubated with HEK293T cell lysate, was able to specifically pull down HSP90α/β (Fig. 1E). Using the same method, we demonstrated specific binding of polyP_100_ with endogenous ATP5A, the alpha subunit of the F1 complex of mitochondrial FoF1 ATP synthase (Fig. 1F), which was present in our polyP interatcome (Table S2). We could also validate the binding of polyP to overexpressed mitochondrial proteins GLUD1 (glutamate dehydrogenase 1) and MICU2 (calcium uptake protein 2) (Fig. 1G, H; Table S2).

**Figure 1.**
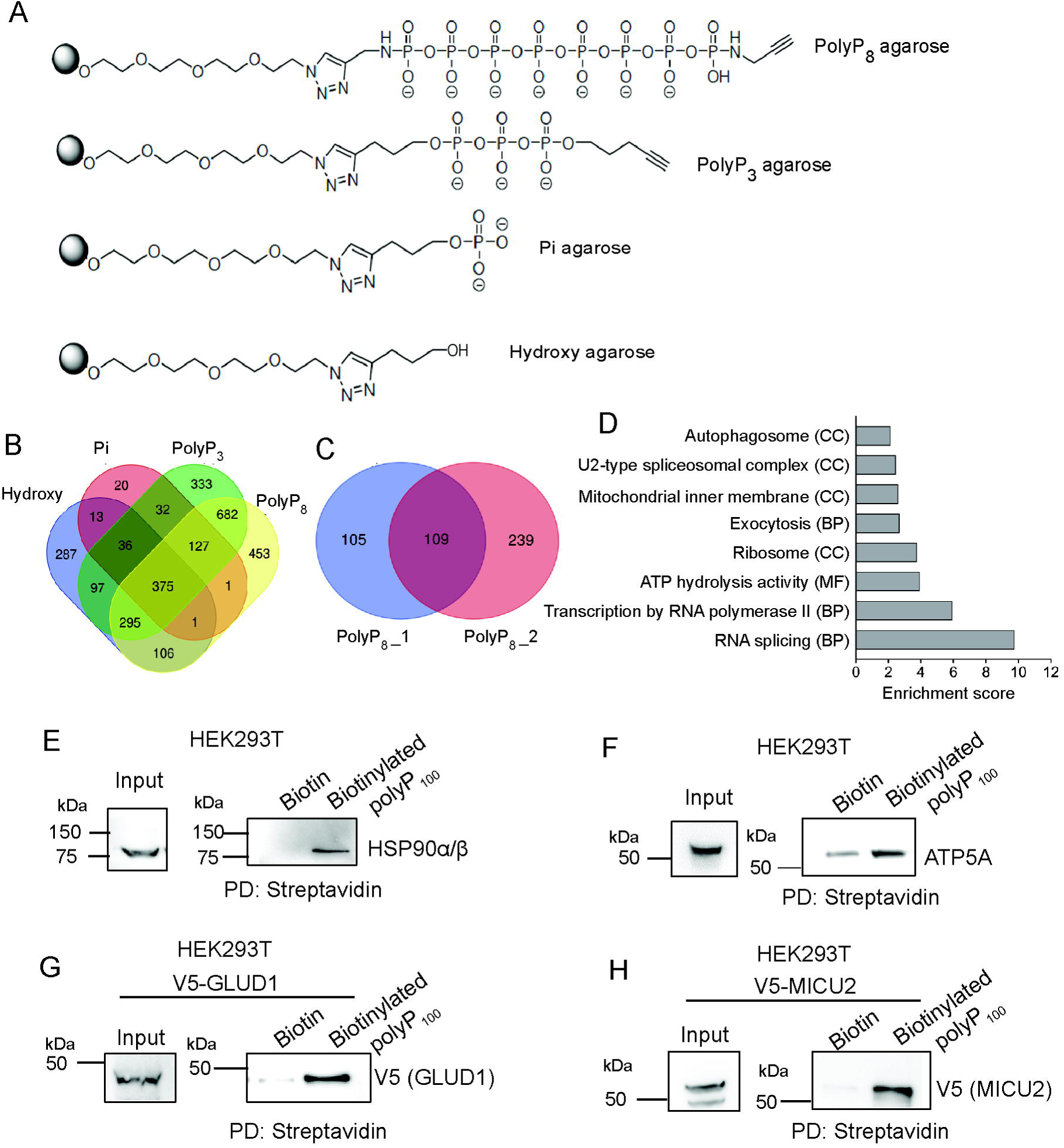
Human polyP interactome analysis and validation: **(A)** Chemical structures of polyP_8_-, polyP_3_-, Pi-, and hydroxy-agarose beads, used to pull-down interacting proteins from HEK293T lysates. **(B)** Venn diagram depicting the number of proteins identified in the pull-down with hydroxy-, Pi-, polyP_3_-, and polyP_8_-agarose beads, combined for two biological replicates in each set (Table S1). **(C)** Venn diagram depicting the number of proteins in each biological replicate for the 453 proteins from (B) that interact exclusively with polyP_8_. **(D)** The total peptide counts for each protein in the test (polyP_8_-agarose) and control (polyP_3_-, Pi-, and hydroxy-agarose) pull-down samples were analyzed using the CRAPome tool (Table S2); proteins that show a fold change (FC-B) score of ≥ 1.5 were subjected to Functional Annotation Clustering of Gene Ontology (GO) terms using the DAVID tool (Table S3). The graph represents the Cellular Component (CC), Biological Process (BP) and Molecular Function (MF) GO terms in clusters with a group enrichment score ≥ 2 (P≤0.01). **(E-F)** Co-precipitation of endogenous HSP90α/β (E) and ATP5A (F) with biotinylated polyP_100_. HEK293T lysate was incubated with either biotin or biotinylated polyP_100_ immobilized on streptavidin beads and probed to detect endogenous HSP90A and ATP5A. **(G-H)** Co-precipitation of GLUD1 and MICU2 with biotinylated polyP_100_. V5-tagged GLUD1 or MICU2 were expressed in HEK293T, pulled down using either biotin or biotinylated polyP_100_ immobilized on streptavidin beads, and probed to detect the V5 tag.

### Biochemical measurement of polyP using DAPI fluorescence

Several methods have been developed for the isolation and measurement of polyP [29–32]. We chose the phenol extraction and ethanol precipitation method, without prior RNAse treatment, which has been shown to yield intact polyP from mammalian cells [30, 33]. The most commonly used method for measurement of polyP involves the degradation of polyP using recombinant *S. cerevisiae* exopolyphosphatase (*Sc*PPX), followed by colorimetric estimation of the released Pi using ammonium molybdate and malachite green [30, 32]. The fluorescent DNA binding dye DAPI, which also binds polyP (Ex_max_ 415 nm, Em_max_ 550 nm), has been used to detect polyP in polyacrylamide gels [34]. To compare these polyP detection methods, we measured different amounts of synthetic polyP of average chain length 20 (polyP_20_), using either DAPI or malachite green, and noted that the polyP-DAPI fluorescence method provided a better dynamic range for polyP estimation compared with the colorimetric method (Fig. S1A, B). It is known that DAPI can also bind RNA with an emission spectrum overlapping with DAPI-polyP [30, 35]. We confirmed that when excited at 415 nm, the emission spectra of DAPI-polyP and DAPI-RNA overlap, with both spectra showing maximum fluorescence in the range of 540-550 nm (Fig. S1C). As our method to isolate polyP co-precipitates cellular RNA, we developed a polyP estimation assay measuring differential DAPI fluorescence in precipitates that are untreated or treated with *Sc*PPX to degrade polyP (Fig. S1D). In a phenol-extracted ethanol precipitate from *S. cerevisiae* (strain BY4741), DAPI fluorescence corresponds to the amount of RNA and polyP in untreated samples, whereas in *Sc*PPX treated samples the remaining fluorescence corresponds to the RNA content in the extract (Fig S1E). The fluorescence values obtained from untreated and *Sc*PPX-treated samples were interpolated on a standard curve of DAPI fluorescence *vs* PolyP_20_ (Fig. S1F). The interpolated value from the *Sc*PPX-treated sample was subtracted from that of the untreated sample, and the difference corresponded to the amount of polyP present in the extract. Employing this method, we quantified polyP levels in *S. cerevisiae* BY4741 strain, and observed values similar to those reported earlier using the malachite green method [36], thereby confirming the reliability of our assay. The validity of our method was further confirmed when the absence of polyP was demonstrated in *vtc1*Δ and *kcs1*Δ strains that lack the yeast polyP synthase and 5-InsP_7_ synthesizing enzyme respectively (Fig. S1G).

### Distribution of polyP in mammalian cells and tissues

Using the DAPI fluorescence assay, we performed polyP quantification in various mammalian cell lines and mouse tissues. PolyP levels were in the range of 15-25 nmoles (in Pi terms)/mg protein in immortalized and tumour-derived cell lines of human and mouse origin (Fig. 2A). PolyP was also detectable by the DAPI fluorescence method in mouse tissues, with highest levels in the testis, and lower levels in mouse liver, brain and kidney (Fig. 2B). Estimation of polyP in different subcellular compartments isolated from cell lines and mouse liver (Fig. S2A-C) revealed that mitochondria contain a higher concentration of polyP compared with the nucleus and post-mitochondrial supernatant (i.e. cytoplasm and other organelles) (Fig. 2C-E). Our data confirms previous reports of the presence of polyP in mitochondria [10, 12, 17, 37], and corroborates an earlier study demonstrating the distribution of polyP in mammalian cell lines, tissues and sub-cellular compartments [4].

**Figure 2.**
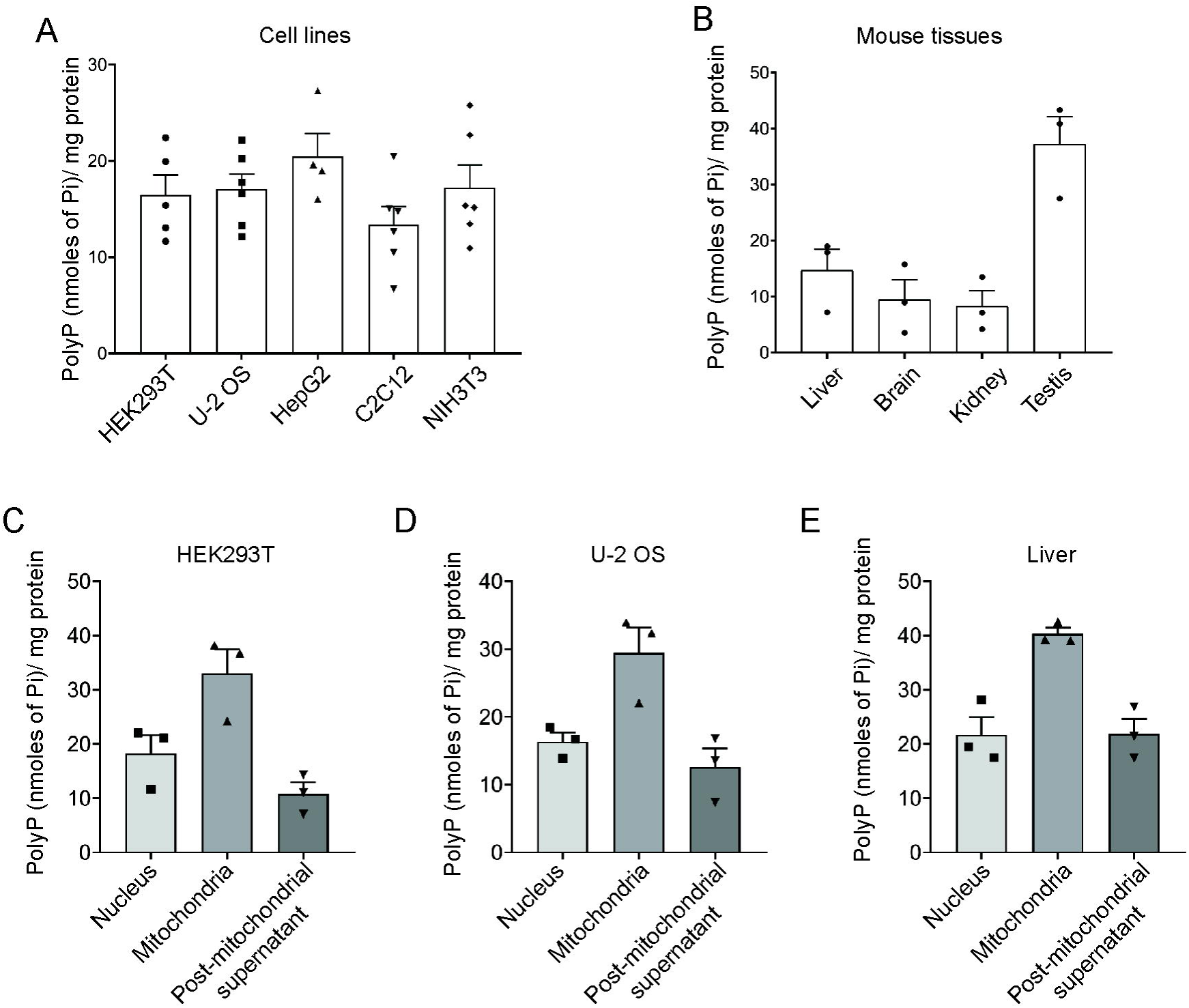
Cellular and subcellular distribution of polyP in mammals. **(A)** Bar graphs depicting the polyP levels (mean ± S.E.M.) in various mammalian cell lines: HEK293T (immortalized human embryonic kidney cell line expressing SV40 large T antigen) (N=5); U-2 OS (human osteosarcoma-derived cell line) (N=6); HepG2 (human hepatocellular carcinoma-derived cell line) (N=4); C2C12 (immortalized mouse myoblast cell line) (N=6); NIH3T3 (immortalized mouse embryonic fibroblast cell line) (N=6). **(B)** Bar graphs representing polyP levels (mean ± S.E.M.) in various tissues including liver, brain, kidney, and testis, isolated from C57BL/6 mice (N=3). **(C-E)** Bar graphs showing the subcellular distribution of polyP in mammalian cells. Levels of polyP (mean ± S.E.M.) in nuclear, mitochondrial, and post-mitochondrial fractions from HEK293T cells (C), U-2 OS cells (D), and mouse liver tissue (E). PolyP levels, expressed in terms of Pi units, were normalized to the total amount of protein in the cell or tissue homogenate used for polyP extraction (N=3).

### Mitochondrial activity is required for polyP synthesis

Previous studies on mitochondrial polyP synthesis have shown that isolated mitochondria incubated with substrates of either complex I or II of the electron transport chain (ETC) show polyP synthesis in the presence of Pi [16, 17]. It has also been shown that treatment of isolated mitochondria with ETC inhibitors blocks polyP synthesis. However, the relationship between mitochondrial respiration and polyP levels has never been studied. Thus, we systematically investigated whether mitochondrial polyP levels are affected by mitochondrial activity. When cultured in high glucose (25 mM), highly proliferative cells, such as cancer cells prefer glycolysis over oxidative phosphorylation for ATP production, even in the presence of oxygen [38]. In low glucose (5 mM), these cells undergo a metabolic shift from glycolysis to fatty acid oxidation and rely on mitochondrial respiration for ATP synthesis [39, 40]. When glucose in the medium is replaced with galactose (10 mM) in the presence of glutamine (2 mM), mitochondrial respiration is stimulated and cells in culture rely solely on oxidative phosphorylation for ATP generation [40, 41]. We observed that when U-2 OS cells were cultured in either low glucose or galactose-containing DMEM for 12 h, mitochondrial polyP levels increased by 2-3 fold compared with cells cultured in high glucose (Fig. 3A). To further examine the effect of mitochondrial respiration on polyP, we treated cells grown in low glucose with mitochondrial inhibitors. Antimycin A, an inhibitor of Cytochrome c reductase (complex III), impedes the flow of electrons in the electron transport chain (ETC) to disrupt the proton gradient across the inner mitochondrial membrane [42]. Treatment of U-2 OS cells with antimycin A resulted in a nearly 50% decrease in mitochondrial polyP levels (Fig. 3B). Treatment with the protonophore FCCP (carbonyl cyanide-p-trifluoromethoxyphenylhydrazone), which collapses the mitochondrial membrane potential to uncouple ATP synthesis from the ETC [43], led to a 20% decrease in mitochondrial polyP (Fig. 3B). Oligomycin inhibits the mitochondrial FoF1 ATP synthase (complex V) by blocking proton entry through subunit C of the Fo complex, in the process increasing mitochondrial membrane potential [44]. In the presence of oligomycin, mitochondrial polyP levels fell by approximately 20% (Fig. 3B). Our data therefore confirm that maintenance of polyP levels in the mitochondria requires an intact proton gradient across the inner mitochondrial membrane, and a functional FoF1 ATP synthase [16, 17].

**Figure 3.**
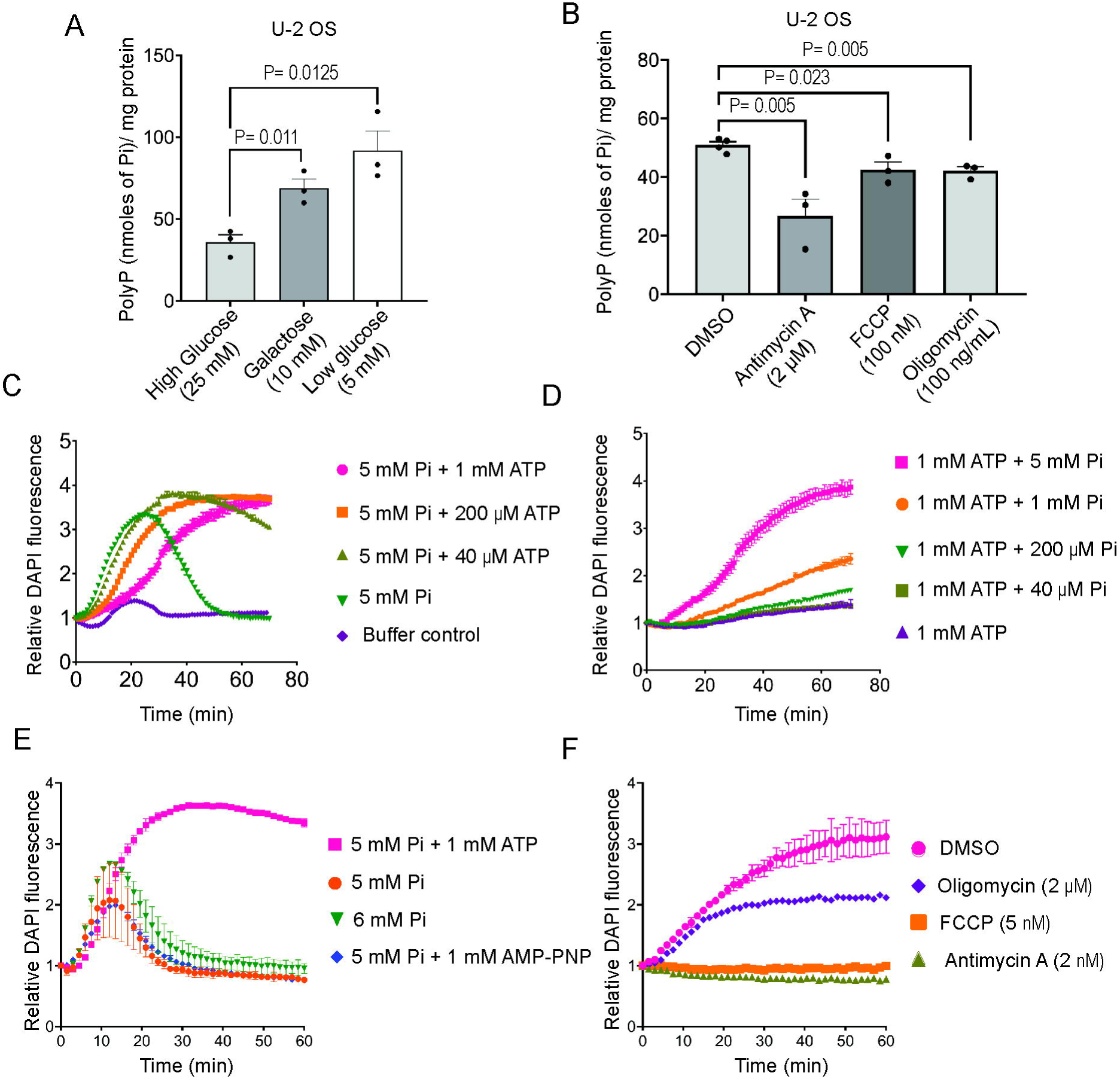
Effect of mitochondrial activity on polyP synthesis. **(A)** Bar graph depicting mitochondrial polyP levels (mean ± S.E.M.) in U-2 OS cells cultured in high-glucose (25 mM), galactose (10 mM), or low-glucose (5 mM) containing DMEM. P values were determined using a two-tailed unpaired Student’s *t*-test (N=3) **(B)** Bar graph representing mitochondrial polyP levels in U-2 OS cells treated for 3 h with Antimycin A (2 µM; complex III inhibitor), FCCP (100 nM; protonophore and uncoupler of oxidative phosphorylation), Oligomycin (100 ng/mL; complex V inhibitor), or DMSO (vehicle control). P values were determined using a two-tailed unpaired Student’s *t*-test (N=3). **(C-F)** Line graphs representing the fold change in DAPI fluorescence over time as a measure of polyP synthesis in isolated mitochondria incubated with glutamate (5 mM), malate (5 mM), and succinate (10 mM), under different conditions as indicated: high Pi (5 mM) with varying concentrations of ATP (C); high ATP (1 mM) with varying concentrations of Pi (D); addition of ATP (1 mM), non-hydrolyzable ATP analog AMP-PNP (1 mM), or Pi (1 mM), along with 5 mM Pi (E); Pi (5 mM) and ATP (1 mM), along with mitochondrial inhibitors at the indicated concentrations, or DMSO as a control (F).

To further understand the molecular details of mitochondrial polyP metabolism, we conducted *ex vivo* polyP synthesis in mitochondria isolated from mouse liver. We first used JC-1 dye to confirm that the membrane potential is intact in isolated mitochondria over a period of 90 min at 30°C (Fig. S2D). We then confirmed that there is no interference from ATP when the DAPI-polyP complex is monitored at Ex_415 nm_/Em_550 nm_ (Fig. S2E). Isolated mitochondria were incubated with substrates of the tricarboxylic acid (TCA) cycle - glutamate (5 mM), malate (5 mM), and succinate (10 mM), that provide NADH and FADH_2_ to complex I and complex II of the ETC respectively. We added varying concentrations of orthophosphate (Pi) and ATP, keeping one or the other constant, and continuously monitored polyP synthesis using DAPI fluorescence. The addition of Pi alone permitted polyP synthesis in isolated mitochondria, but ATP alone did not support polyP synthesis (Fig. 3C, D). In the presence of Pi alone, polyP levels peaked and fell during the period of observation, and the addition of increasing concentrations of ATP led to a delay in reaching peak polyP levels while enhancing polyP stability (Fig. 3C). Increasing the concentration of Pi in the presence of 1 mM ATP (which approximates steady state cellular ATP levels) led to a steady increase in polyP synthesis (Fig. 3D). These data suggest that Pi serves as the primary source of phosphate for polyP production in the mitochondria, whereas ATP enhances mitochondrial polyP synthesis. To understand whether ATP supports polyP synthesis via hydrolysis or regulatory binding, we utilized AMP-PNP, a non-hydrolysable form of ATP. Interestingly, the addition of AMP-PNP did not enhance Pi-driven polyP synthesis in isolated mitochondria (Fig. 3E), indicating that ATP hydrolysis is needed for sustained polyP synthesis. Increasing the concentration of Pi from 5 mM to 6 mM did not have the same impact on polyP synthesis as the addition of 1 mM ATP to 5 mM Pi. Together these data suggest that ATP hydrolysis does not merely act to provide additional Pi equivalents that go towards polyP, but instead drives the mitochondrial polyP synthesis machinery by another, yet to be determined, energy-driven mechanism. Finally, we assessed the impact of mitochondrial inhibitors on *ex vivo* polyP synthesis in the presence of ATP (1 mM) and Pi (5 mM). The inclusion of antimycin A or FCCP led to a complete abrogation of polyP synthesis (Fig. 3F), confirming that an intact mitochondrial membrane potential is pre-requisite for the production of polyP in isolated mitochondria. The addition of increasing doses of oligomycin lowered, but did not block polyP synthesis (Fig. 3F, S2F), suggesting that activity of the FoF1ATP synthase directly or indirectly supports mitochondrial polyP production.

### IP6K1 maintains mitochondrial polyP levels via the synthesis of 5-InsP_7_

In budding yeast, the inositol pyrophosphate 5-InsP_7_ upregulates polyP synthesis via allosteric modulation of the VTC complex, the yeast polyP synthase [18–20]. In mammals, 5-InsP_7_ is synthesised from InsP_6_ by IP6 kinases, of which there are three paralogs – IP6K1/2/3. We have earlier shown that the loss of IP6K1 in mice leads to a reduction in platelet polyP, with a concomitant impairment in blood clotting [21]. To determine whether IP6K1 also influences mitochondrial polyP, we isolated mitochondria from cell lines and liver of mice lacking IP6K1. In HEK293T and U-2 OS cells in which both alleles of IP6K1 have been deleted using CRISPR-Cas9 editing, we observed a 30-40% decrease in mitochondrial polyP levels compared with their respective non-targeted control (NTC) lines (Figs. 4A, B and S3A, B). In mitochondria isolated from the liver of *Ip6k1*^-/-^ mice, we see a 40% decrease in polyP compared with *Ip6k1*^+/+^ mice (Fig. 4C, S3C). Unlike control cells, U-2 OS cells lacking IP6K1 did not show a significant increase in mitochondrial polyP levels when they were grown for 12 h in low glucose compared with high glucose containing medium (Fig. 4D), suggesting that mitochondrial activity may be impaired in the absence of IP6K1.

**Figure 4.**
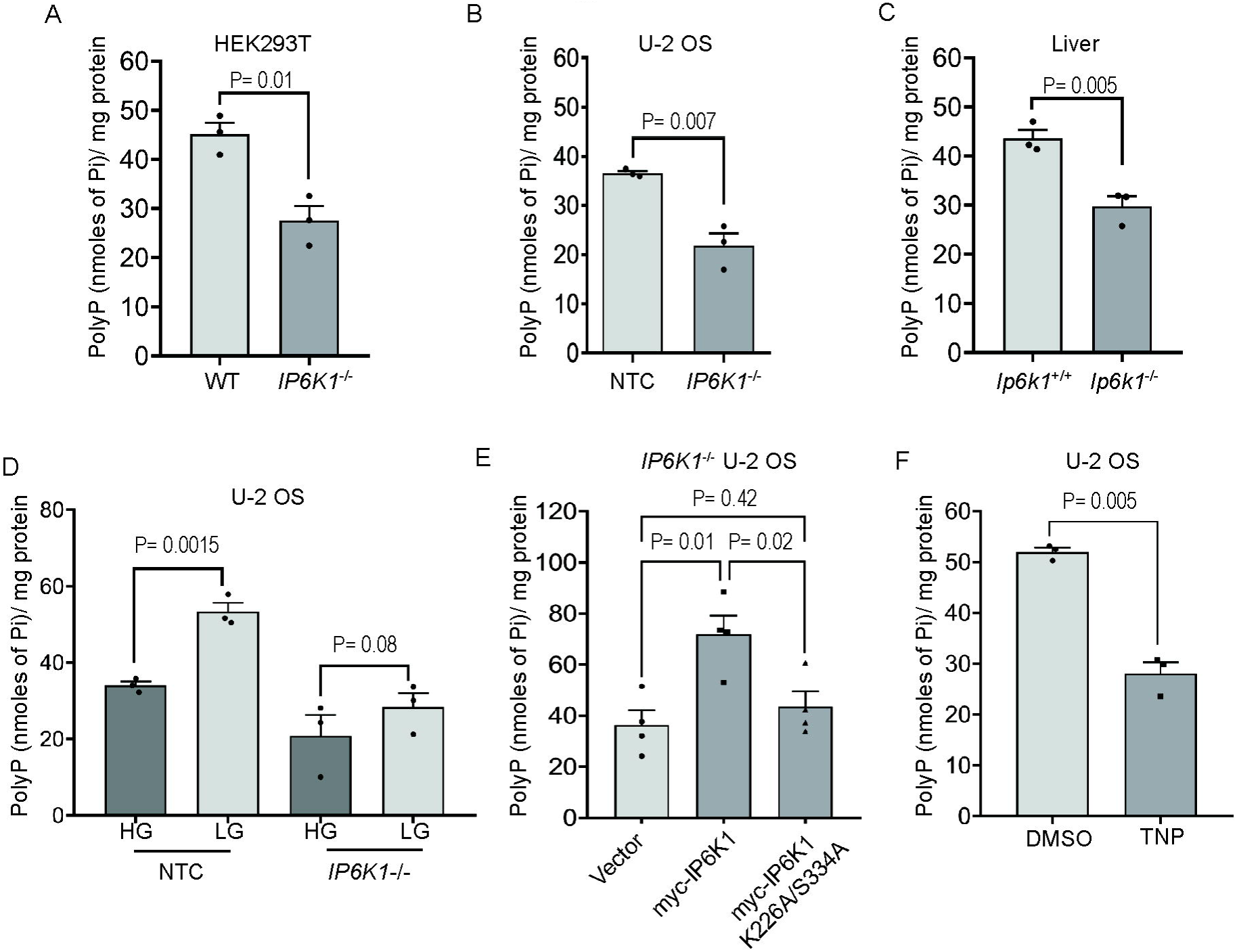
IP6K1 regulates mitochondrial polyP levels. **(A-C)** Bar graphs illustrating the levels of mitochondrial polyP (mean ± S.E.M.) in WT and *IP6K1*^-/-^ HEK293T cells (N=3) (A), non-targeting control (NTC) and *IP6K1*^-/-^ U-2 OS cells (N=3) (B), and *Ip6k1*^+/+^ and *Ip6k1*^-/-^ mouse liver (N=3) (C). **(D)** Bar graph representing mitochondrial polyP levels (mean ± S.E.M.) in NTC and *IP6K1*^-/-^ U-2 OS cells cultured in high glucose (25 mM; HG) or low glucose (5 mM; LG) medium for 12 h (N=3). **(E)** Bar graph representing mitochondrial polyP levels in *IP6K1*^-/-^ U-2 OS cells stably expressing myc-tagged active or catalytically inactive (K226A/S334A) mouse IP6K1 (N=3). **(F)** Bar graph representing mitochondrial polyP levels in U-2 OS cells treated with 10 µM TNP (IP6K inhibitor). DMSO was used as vehicle control (N=3). All data were analysed for P values using a two-tailed unpaired Student’s *t*-test.

To determine whether IP6K1 regulates mitochondrial polyP levels via the synthesis of 5-InsP_7_ or by a mechanism independent of its catalytic activity, we generated *IP6K1^-/-^* U-2 OS derived cell lines that stably express myc-tagged mouse IP6K1 that is active or catalytically inactive (K226A/S334A) (Fig. S3D). We confirmed that the loss of IP6K1 leads to a reduction in cellular levels of 5-InsP_7_ (Fig. S3E), and that the expression of active IP6K1 but not mutant IP6K1 can restore 5-InsP_7_ in *IP6K1^-/-^* U-2 OS cells (Fig. S3F). The expression of active IP6K1 increased polyP levels in *IP6K1^-/-^* U-2 OS cells cultured in low glucose, whereas mutant IP6K1 did not show a change in polyP levels compared with control cells expressing vector alone (Fig. 4E), suggesting that 5-InsP_7_ supports the maintenance of mitochondrial polyP. Treatment of U-2 OS cells cultured in low glucose with the pan-IP6K inhibitor TNP (10 µM for 16 h) led to a ∼50% decrease in polyP, confirming that 5-InsP_7_ synthesis is required to maintain mitochondrial polyP levels.

### IP6K1 regulates mitochondrial activity

Having established that IP6K1 supports mitochondrial polyP via the synthesis of 5-InsP_7_, and observed that the maintenance of mitochondrial membrane potential is needed for polyP synthesis, we probed whether IP6K1 regulates mitochondrial function. We first checked whether IP6K1 localises to the mitochondria by co-staining U-2 OS cells with mitotracker red, a dye that marks the mitochondrial matrix, and an antibody to detect endogenous IP6K1. We saw IP6K1 staining in the nucleus and cytoplasm as reported earlier [45], but failed to see any co-localisation of IP6K1 with the mitochondrial marker (Fig. S4A). We were also unable to detect any enrichment of IP6K1 in the mitochondrial fraction in U-2 OS cells (Fig. S4B). An earlier study has shown that *Ip6k1*^-/-^ mouse embryonic fibroblasts display reduced mitochondrial respiration compared with their wild type counterparts [46]. We used the Seahorse assay system [47] (Fig S4C) to assess cellular respiration in NTC and *IP6K1^-/-^* U-2 OS cells. In control cells assayed in low glucose, we observed a significant increase in basal oxygen consumption rate (OCR) compared with cells in high glucose medium (Fig. 5A, C) reflecting an increase in mitochondrial respiration in glucose deprived conditions. Injection of oligomycin into the assay system inhibited ATP-linked mitochondrial respiration, and the remaining oxygen consumption, associated with proton leak across the inner mitochondrial membrane, was not substantially different in low and high glucose medium (Fig. 5A), implying that the shift to low glucose increases ATP-linked respiration. The addition of FCCP, which allows the ETC to function at its maximum capacity, led to ATP synthesis-independent maximal respiration and restored the difference in OCR observed between low and high glucose conditions (Fig. 5A, C). The addition of rotenone (an inhibitor of Complex I) along with antimycin A abrogates mitochondrial respiration, and the remaining non-mitochondrial respiration was not significantly dependent on glucose levels in the medium (Fig. 5A). In *IP6K1^-/-^* U-2 OS cells, basal respiration was lower than that observed in NTC cells, and remained unchanged regardless of the glucose concentration in the media (Fig. 5B, C). Addition of oligomycin confirmed that ATP-linked respiration is indeed lower in *IP6K1^-/-^* U-2 OS cells. Subsequent addition of FCCP failed to show any effect of low glucose on maximal respiration in

**Figure 5:**
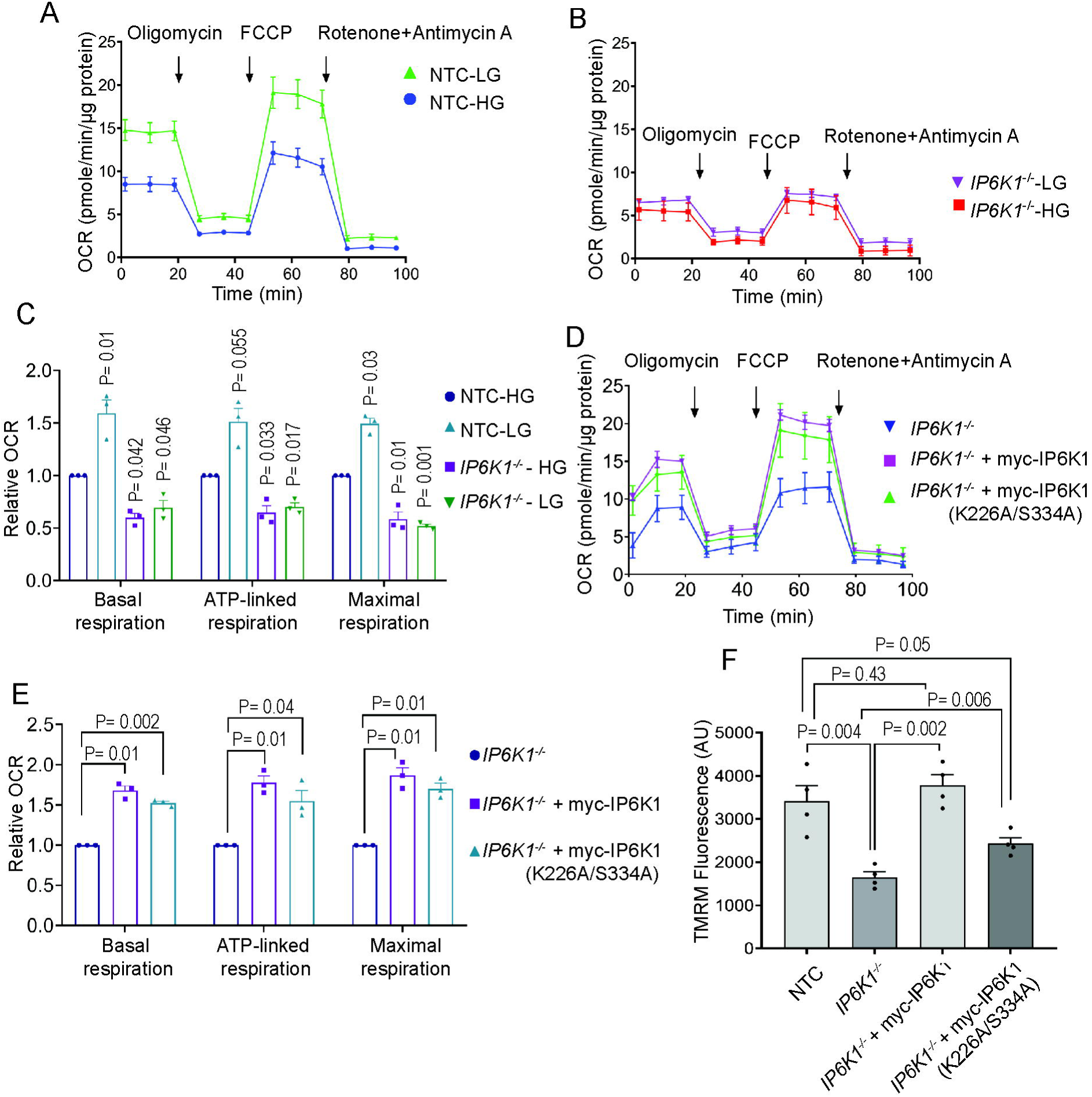
Mitochondrial respiration is regulated by IP6K1: **(A, B)** Oxygen consumption rate (OCR) traces, expressed as pmoles of O_2_/min/µg protein for NTC (A) and *IP6K1*^-/-^ (B) U-2 OS cells cultured in high glucose (HG) or low glucose (LG) medium. Arrows indicate the time points at which Oligomycin, FCCP, or Antimycin A and Rotenone were added during the assay. The OCR profiles are representative of three independent experiments. **(C)** Bar graph representing relative fold change in OCR values of the indicated U-2 OS cell lines in HG or LG medium, normalised to OCR values of NTC cells in HG medium. Data (mean ± S.E.M.) were analyzed using a one-sample *t*-test (N=3). **(D)** OCR traces for *IP6K1*^-/-^ and *IP6K1*^-/-^ U-2 OS cells stably expressing active or catalytically inactive (K226A/S334A) mouse IP6K1. The OCR profile is representative of three independent experiments. **(E)** Bar graph representing relative fold change in OCR values in LG medium for *IP6K1*^-/-^ U-2 OS cells stably expressing active or catalytically inactive mouse IP6K1, normalised to *IP6K1*^-/-^ U-2 OS cells. Data (mean ± S.E.M.) were analyzed using a one-sample t-test. (N=3). (F) Bar graph depicting TMRM fluorescence values (mean ± S.E.M.) in NTC, *IP6K1*^-/-^ and *IP6K1*^-/-^ stably expressing active or catalytically inactive mouse IP6K1. P values were determined using a two-tailed unpaired Student’s *t*-test (N=3).

*IP6K1^-/-^*U-2 OS cells, which remained lower in IP6K1 depleted cells compared with non-targeted control cells.

Next, we examined whether IP6K1 supports mitochondrial respiration via the synthesis of 5-InsP_7_ or by a catalytic activity-independent mechanism. We monitored OCR parameters in *IP6K1^-/-^* U- 2 OS cells expressing active or mutant IP6K1 alongside the parent *IP6K1^-/-^*U-2 OS cell line. Unlike the effect on mitochondrial polyP levels (Fig. 4E), expression of either active or inactive IP6K1 could restore mitochondrial respiration, although mutant IP6K1 showed marginally lower OCR parameters than active IP6K1 but this difference was not statistically significant (Fig. 5D, E). As mitochondrial respiration is dependent on the proton gradient across the inner mitochondrial membrane, we loaded cells with the fluorescent reporter tetramethylrhodamine methyl ester (TMRM), which accumulates in healthy mitochondria with a functional membrane potential. TMRM fluorescence monitored by flow cytometry was significantly reduced in *IP6K1^-/-^* U-2 OS cells compared with control cells (Fig. 5F) revealing that IP6K1 is needed to maintain the electrical gradient across the inner mitochondrial membrane, and suggesting that impaired respiration in IP6K1 depleted cells may be attributed to a reduction in the mitochondrial membrane potential. Expression of active IP6K1 restored the membrane potential in *IP6K1^-/-^* U-2 OS cells to the level of control cells, whereas mutant IP6K1 restored the membrane potential only partially. Together, these data reveal a dual role for IP6K1 in the maintenance of healthy mitochondria – IP6K1 acts via the production of 5-InsP_7_ to maintain the proton gradient across the mitochondrial membrane, while acting additionally at the protein level, independent of its catalytic activity, to support mitochondrial respiration.

### IP6K1 maintains mitochondrial polyP synthesis

So far we have seen that polyP synthesis in isolated mitochondria requires an intact proton gradient across the inner mitochondrial membrane and that the loss of IP6K1 negatively impacts the membrane potential and respiration in the mitochondria. IP6K1 depletion is therefore likely to impair mitochondrial polyP synthesis. To test this, we monitored polyP synthesis activity in mitochondria isolated from *Ip6k1^+/+^* and *Ip6k1^-/-^* mouse liver. *Ex vivo* polyP synthesis was significantly lower but not abrogated in mitochondria isolated from the liver of mice lacking IP6K1 (Fig. 6A). This observation was recapitulated in mitochondria isolated from *IP6K1^-/-^* U-2 OS cells, which showed minimal polyP synthesis compared with control cells (Fig. 6B). The expression of active IP6K1 was able to restore polyP synthesis *IP6K1^-/-^*U-2 OS cells whereas inactive IP6K1 did not rescue this defect. These *ex vivo* polyP synthesis assays in isolated mitochondria corroborate our observation of reduced mitochondrial polyP levels in 5-InsP_7_ depleted cells (Fig. 4E, F), and point to a direct or indirect role for 5-InsP_7_ in the maintenance of mitochondrial polyP.

**Figure 6.**
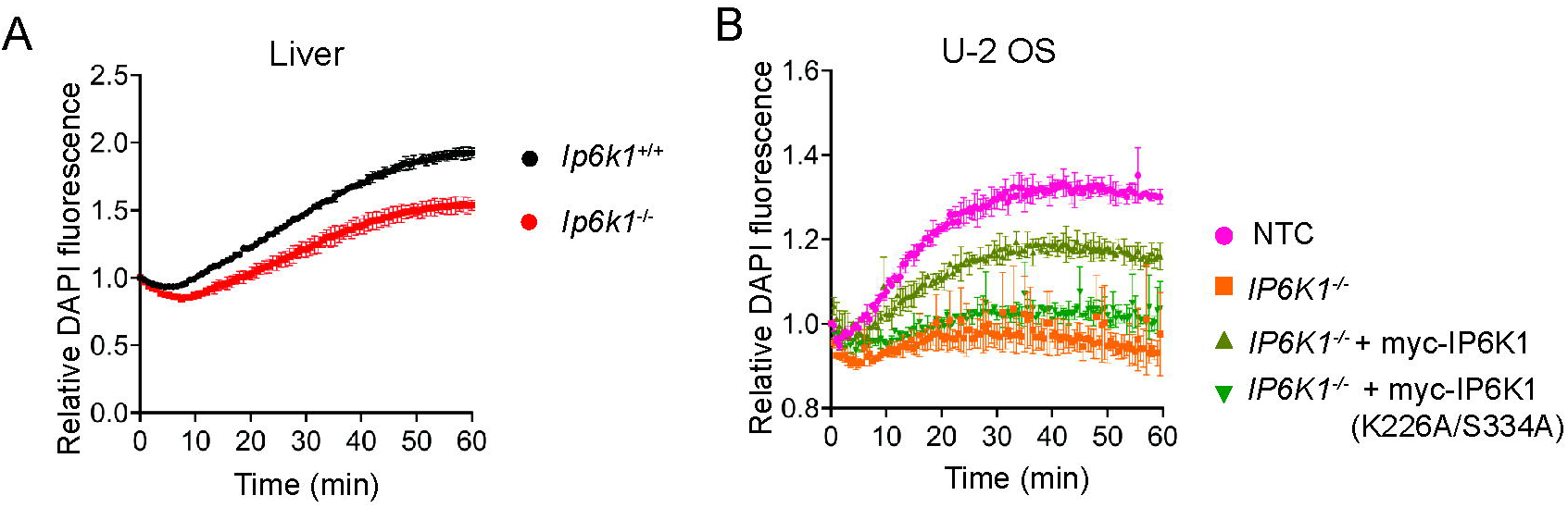
IP6K1 regulates mitochondrial polyP synthesis. **(A, B)** Line graphs representing the fold change in DAPI fluorescence over time as a measure of polyP synthesis in isolated mitochondria from *Ip6k1*^+/+^ and *Ip6k1*^-/-^ mouse liver (N=4) (A), and NTC, *IP6K1*^-/-^, and *IP6K1*^-/-^ U-2 OS cells stably expressing myc-tagged active or catalytically inactive mouse IP6K1 (K226A/S334A) (N=4) (B), incubated with glutamate (5 mM), malate (5 mM), succinate (10 mM), Pi (5 mM), and ATP (1 mM).

## DISCUSSION

PolyP, despite its simplicity of structure and ubiquitous occurrence across living organisms, remains an understudied biopolymer. Mammalian polyP has been especially difficult to study due to its low abundance and poorly understood biochemistry. Here, we have shed light on mechanisms regulating the synthesis of polyP in mammalian mitochondria. We show that mitochondrial polyP synthesis relies on Pi as a substrate and that ATP hydrolysis augments polyP synthesis in isolated mitochondria. Our data confirm that the potential gradient across the inner mitochondrial membrane and activity of the FoF1 ATP synthase are required for mitochondrial polyP synthesis. Additionally, our work has unveiled a role for IP6K1, and its product - the inositol pyrophosphate 5-InsP_7_, in modulating mitochondrial polyP synthesis. Mouse liver and cell lines depleted for IP6K1 and 5-InsP_7_ have reduced levels of mitochondrial polyP, and mitochondria isolated from these cells and tissue show impaired polyP synthesis. Cells lacking IP6K1 display compromised mitochondrial health, as seen from their reduced respiration and mitochondrial membrane potential. We see that IP6K1 supports mitochondrial function both by the synthesis of 5-InsP_7_, and also via a mechanism that is independent of its catalytic activity.

Our data on real time polyP synthesis measured by DAPI fluorescence in mitochondria isolated from mouse liver corroborate similar studies in mitochondria obtained from the liver of Sprague-Dawley rats [16, 17]. As reported earlier, we see that mitochondrial polyP synthesis requires substrates of complex I and complex II of the ETC, and an intact potential gradient across the inner mitochondrial membrane. In contrast to earlier reports, we see a reduction, but not an abrogation of mitochondrial polyP synthesis in the presence of oligomycin at 2 μg/mL; in our hands, raising the concentration of oligomycin up to 20 μg/mL reduced, but did not block polyP synthesis. One critical point of agreement between our data and previous reports is the requirement of Pi to support mitochondrial polyP synthesis [16, 17]. On the other hand, a major point of divergence is that an earlier study reported no effect of ATP on polyP synthesis in rat liver mitochondria, whereas we observed that the addition of ATP potentiates and sustains Pi-driven polyP synthesis in isolated mouse liver mitochondria. By showing that neither the non-hydrolysable ATP analog AMP-PNP, nor an equivalent concentration of additional Pi, recapitulate the effect of ATP on promoting Pi-driven polyP synthesis, our data suggests that ATP hydrolysis promotes mitochondrial polyP synthesis by a currently unknown mechanism. Overall, our data support the contention that the FoF1 ATP synthase may also function as a mitochondrial polyP synthase [17], with the caveat that mitochondrial polyP synthesis in the presence of oligomycin may be mediated by another enzyme or by a fraction of FoF1 synthase that is resistant to oligomycin. If FoF1 indeed acts as a polyP synthase, it raises intriguing possibilities regarding a metabolic switch between ATP and polyP synthesis, potentially regulated by ADP availability and prevailing cellular ATP levels. The functional significance of mitochondrial polyP may lie in its ability to serve as a readily accessible reservoir of Pi within the organelle [2]. FoF1 ATPase has also been shown to hydrolyse polyP [17], and may be responsible for the release of polyP-derived Pi units into the mitochondrial matrix. The spatiotemporal organization and regulation of pools of FoF1 ATP synthase in the mitochondria responsible for synthesis or hydrolysis of ATP or polyP would be an intriguing area of future study.

Despite the divergence in the enzymatic routes for their synthesis, our study brings out two points of similarity between polyP synthesis in mammalian mitochondria and in the vacuoles of budding yeast. The first is the essentiality of an intact proton gradient across the organelle membrane within which polyP accumulates, and the second is a regulatory role for the inositol pyrophosphate 5-InsP_7_. It remains to be determined whether polyP synthesis in mammalian lysosome related granules such as platelet dense granules also relies on a similar potential gradient, but it has been shown previously that depletion of IP6K1 impairs the accumulation of polyP in platelet granules in mice [21]. This conserved theme of 5-InsP_7_ acting as a regulator of organellar polyP accumulation is in sync with the well known role of 5-InsP_7_ as an energy sensor. Owing to the low affinity of IP6Ks for ATP (Km ∼1 mM), cellular levels of 5-InsP_7_ are acutely sensitive to fluctuations in cellular ATP concentrations [48–51]. Budding yeast devoid of 5-InsP_7_ have dysfunctional mitochondria but, paradoxically, contain four times as much ATP because of increased glycolysis [46]. Similarly, mouse embryonic fibroblasts derived from *Ip6k1^-/-^* mice have reduced mitochondrial activity and membrane potential, suggesting a regulatory role of IP6K1 in mitochondrial metabolism [46]. We observed that in U-2 OS cells, IP6K1 regulates mitochondrial respiration and membrane potential by mechanisms that are independent of or dependent on its ability to catalyse the synthesis of 5-InsP_7_. While the precise molecular mechanisms remain to be elucidated, it is plausible that 5-InsP_7_ exerts its effects on mitochondrial membrane potential, and consequently on mitochondrial polyP synthesis, through binding to or pyrophosphorylation of proteins in the nucleus or cytoplasm. Additionally, IP6K1, independent of its catalytic activity, may influence mitochondrial respiration through regulatory protein-protein interactions. Further research is needed to fully unravel the intricate interplay between polyP metabolism, mitochondrial membrane potential, and the IP6K1/5-InsP_7_ signalling pathway in maintaining mitochondrial homeostasis.

## MATERIALS AND METHODS

### Reagents

All chemicals were procured from Sigma-Aldrich, unless specified otherwise. The primary antibodies used in the present study for immunoblotting, along with antibody dilution for each application, and supplier (including catalogue number), were as follows: anti-HSP90 (Abcam, Ab13492, 1:2000); anti-ATP5A (Abcam, Ab14748, 1:5000); anti-V5 tag (Thermo Fisher Scientific, R960-25, 1:5000); anti-LaminB1 (Abcam, Ab1160828, 1:5000); anti-GAPDH (Sigma-Aldrich, G8795, IB, 1:10,000); anti-IP6K1 (Sigma-Aldrich, HPA040825; IF, 1:300; Genetex, GTX103949; IB, 1:3,000). PVDF membrane for protein transfer, streptavidin-sepharose beads, and ECL prime chemiluminescence substrate were procured from GE Healthcare. *myo*-2-[^3^H] inositol (15-20 Ci/mmol) (ART 0116B) was procured from American Radiolabeled Chemicals. Ultima-Flo AP (6013599) was purchased from Perkin-Elmer.

### Cell lines and transfections

All cell lines were maintained in a humidified incubator with 5% CO_2_ at 37°C, using Dulbecco’s modified Eagle’s medium (DMEM) supplemented with 10% fetal bovine serum (FBS), 1 mM L-glutamine, 100 U/ml penicillin, and 100 µg/ml streptomycin. *IP6K1^-/-^* HEK293T cells were generated using non-targeted *sg*RNA and IP6K1-directed *sg*RNA, respectively, as described previously [52]. NTC and *IP6K1^-/-^*U-2 OS cell lines were established through a CRISPR-Cas9-mediated knockout strategy. Specifically, non-targeted sgRNA or IP6K1-targeted sgRNA sequences were designed by TransOMIC Technologies and cloned into the pCLIP-ALL-EFS-Blasticidin destination vector. Lentiviral particles were produced in HEK293T cells by co-transfecting with VSV-G, psPAX2, and the pCLIP-ALL-EFS-Blasticidin-sgRNA constructs. 48 hours post-transfection, the culture supernatants were collected and filtered through a 0.45 μm filter to isolate the viral particles, which were then used to infect U-2 OS WT cells seeded in 60 mm dishes. Selection of the transduced cells was carried out by treating the cells with 5 μg/mL blasticidin for 15 days. Following selection, single cells were plated in 96-well plates through serial dilution in conditioned media. The surviving colonies were genotyped for frameshift mutations, expanded further, and screened for the IP6K1 knockout through western blot analysis. *IP6K1^-/-^* U-2 OS cell lines, stably expressing active or catalytically inactive myc-tagged IP6K1 (K226A/S334A) IP6K1 (mutant) were generated using retroviral transduction as described earlier [53, 54]. Briefly, the BOSC23 cells were transfected with retroviral vectors pCX-Neo plasmid containing myc-tagged mouse IP6K1 that is active or catalytically inactive (K226A/S334A), and pClAmpho retroviral packaging (NBP2-29541) plasmids. 48 h post-transfection, retrovirus particles were collected by filtering through a 0.45 μm filter, and were used to transduce U-2 OS IP6K1^-/-^ cells. 48 h post transduction, transduced cells were selected using G418 (0.7 mg/mL Sigma-Aldrich Cat no A1720) for 7 days. The selected pool of transduced cells was further maintained in 0.2 mg/mL G418. pLex 983-Micu2 was a gift from Vamsi Mootha (Addgene plasmid #50372). cDNA of GLUD1 was subcloned in pLX304-V5. pLX304 was a gift from David Root (Addgene plasmid # 25890) using gateway cloning method. For transfection, polyethylenimine (PEI) (Polysciences, 23966) was used at a ratio of 1:3 (DNA: PEI). All plasmids used for transfection were purified using the Plasmid Midi kit (Qiagen). Cells were harvested 36-48 h post-transfection for further analyses.

### Mice

All animal experiments were approved by the Institutional Animal Ethics Committee (Protocol number EAF/RB/01/2025), and were performed in compliance with guidelines provided by the Committee for the Control and Supervision of Experiments on Animals, Government of India (2035/GO/RBi/S/2018/CCSEA). *Ip6k1*^+/+^ and *Ip6k1*^-/-^ mice (*Mus musculus*, strain C57BL/6) used for this study were housed in the Experimental Animal Facility at the Centre for DNA Fingerprinting and Diagnostics, Hyderabad. *Ip6k1*^+/+^ and *Ip6k1*^-/-^ littermates were produced by breeding *Ip6k1*^+/-^ mice and maintained as previously described [55].

### Procedure for synthesis of polyP-coupled agarose beads

Clickable polyphosphate conjugates were obtained as described in [24, 26, 56]. The procedure for Click-reactions with azide agarose was adapted from [57]. Azide agarose mixture (0.25 mL, 2.5 - 5 µmol) was washed with water (3 × 1.0 mL) and suspended in MeOH:H_2_O (1:1, 188 µL). To this suspension, the alkyne (1.25 µmol, 1.0 eq.) in MeOH:H_2_O (1:1, 25 µL) was added, followed by the addition of CuSO_4_×5 H_2_O (100 µM, 12.5 µL, 1.25 µmol, 1.0 eq.) and sodium ascorbate (100 µm, 62.5 µL, 6.25 µmol, 5.0 eq.). After the addition of H_2_O (80.3 µL) the mixture was rotated overnight at RT. The suspension was washed with water (3 × 1.0 mL) and the remaining azide functionalities were blocked by repeating the above procedure. Instead of the addition of the modified alkyne, propargylamine (32.0 µL, 500 µmol, 400 eq.) was added. The suspension was rotated for 3 h and was then washed with water (3 × 1.0 mL). The synthesis of agarose-PEG3 labeled phosphate conjugate (Pi-agarose), was achieved using pent-4-yn-1-yl phosphate (418 mg, 1.26 mmol) according to the general procedure. As no starting material was left after the reaction, the loading of the beads is ca. 25 - 50%. Agarose PEG3 labeled triphosphate conjugate (polyP_3_- agarose) was synthesised using bis-pentinyl triphosphate (656 mg, 1.15 mmol), and agarose-PEG3 labeled octaphosphate conjugate (polyP_8_-agarose) was synthesized using bis-propargylamido octaphosphate (1.00 mg, 1.15 mmol), according to the general procedure [26]. As no starting material was left after the reaction in both cases, the loading of the beads is ca. 23 - 46%. The synthesized beads were stored in sodium azide (2% in water) with a total volume of 0.5 mL.

### Mass spectrometry-based identification of the polyP interactome

Synthetic bis-alkyne polyP8 or bis-alkyne polyP3 were prepared as described earlier [26]and conjugated with azide-agarose beads using click chemistry. For identification of proteins interacting with polyP, ∼8 million HEK293T cells were collected and lysed by incubating them in lysis buffer (50 mM HEPES, 150 mM NaCl, 1 mM EDTA, 1 mM DTT, 1 mM NaF, 1X protease inhibitor cocktail, 1X phosphatase inhibitor cocktail) for 45 min at 4 ^°^C followed by centrifugation at 8,000 *g* for 10 min at 4 °C. Cell lysates containing 1 mg protein was added to 100 µl each of polyP8-agarose, polyP3-agarose, Pi-agarose, and hydroxy agarose beads (Fig 1A). The volume was made up to 1 mL with lysis buffer. The beads were placed overnight at 4°C on an end-over-end mixer, washed using lysis buffer, boiled in 2X SDS sample buffer for 5 min, loaded on a 12% SDS polyacrylamide gel, allowed to run into the resolving gel up to 1 cm, and visualized by Coomassie Brilliant Blue staining. All proteins in the sample were excised in one gel slice and sent for mass spectrometry analysis to Taplin Biological Mass Spectrometry Facility at Harvard University, U.S.A. Briefly, gel pieces were subjected to in-gel digestion with sequencing-grade trypsin (Promega), and the extracted peptides were resolved by reverse phase HPLC. Eluted peptides were subjected to electrospray ionization (ESI) and then allowed to enter an LTQ Orbitrap Velos Pro ion-trap mass spectrometer (Thermo Fisher Scientific). Peptides were detected, isolated, and fragmented to produce a tandem mass spectrum of specific fragment ions for each peptide. Peptide sequences (and hence protein identity) were determined by matching protein databases from UniProt (https://www.uniprot.org/taxonomy/10090) with the acquired fragmentation pattern by the software program, Sequest (Thermo Fisher Scientific). Mass spectromery data were submitted to the MassIVE repository, a full member of the Proteome Xchange Consortium. Data can be accessed via the URL <https://massive.ucsd.edu>, with the data set identifier MassIVE MSV000098211. The total peptide number for each protein, which reflects relative protein abundance in the sample, was compared between the test (polyP_8_-agarose) and control (polyP_3_-, Pi- and hydroxy-agarose) samples using the CRAPome tool (https://reprint-apms.org) [27]. Empirical Fold Change Score (FCA and FCB) was used to compare the enrichment of proteins in the bait (polyP_8_-agarose) over user control (PolyP_3_-, Pi-, and hydroxy-agarose) in replicates. A fold change score (FCB) of 1.5 was used as a cut-off score to select genuine interactors of polyP_8_. The selected proteins were examined for Gene Ontology (GO) term enrichment using the Functional Annotation Clustering tool at Database for Annotation, Visualization and Integrated Discovery (DAVID) v6.7 (https://david.ncifcrf.gov) [28].

### Validation of interactome by pull-down assay

To validate the interaction of polyP with proteins identified by mass-spectrometry, we used biotinylated polyP_100_. Streptavidin agarose beads (Cytiva, 17511301) were pre-incubated with an equal volume of 4 µM biotinylated polyP_100_ or biotin (negative control) for 1 h with shaking at 4°C to prepare a slurry of 2 µM polyP_100_ immobilized on beads. HEK293T cells were collected 48 h post transfection (in cases of protein over-expression), lysed for 1 h at 4°C in lysis buffer (50 mM HEPES, 150 mM NaCl, 1 mM EDTA, 0.5 mM NaF, 1X phosphate inhibitor cocktail, 1X protease inhibitor cocktail, and 0.5% of Nonidet P-40). The lysate was incubated for 2 h on an end-over-end mixer at 4°C with 10 µL slurry of biotinylated P_100_ or biotin (negative control) immobilised on streptavidin agarose beads. The beads were washed three times with lysis buffer followed by boiling in 1X Laemmli buffer. The samples were analyzed by immunoblotting, and UVITEC Alliance Q9 documentation system or GE Image-Quant LAS 500 imager were used for chemiluminescence detection.

### Subcellular fractionation

#### For mammalian cells

Approximately 25 million cells were harvested at 80-90% confluency and resuspended in hypotonic buffer (10 mM NaCl, 1.5 mM MgCl_2_, 10 mM Tris-HCl pH 7.5, 10 mM NaF, 1X protease inhibitor cocktail, and 1X phosphatase inhibitor cocktail). After incubating on ice for 20 min, the cells were homogenized using a Dounce homogenizer with 20 strokes of Pestle A followed by 20 strokes of Pestle B (repeated twice). The resulting homogenate was mixed with an equal volume of 2X MS homogenization buffer (1X: 210 mM mannitol, 70 mM sucrose, 5 mM Tris-HCl pH 7.5, and 1 mM EGTA). This homogenate was then subjected to differential centrifugation to isolate the nuclear pellet, mitochondrial pellet, and post-mitochondrial supernatant. The total homogenate was first centrifuged at 1,500 *g* for 15 min to remove the nuclear fraction. The resulting supernatant was then centrifuged at 8,000 *g* for 20 min to pellet mitochondria, which were subsequently resuspended in 1X MS homogenization buffer, and the post-mitochondrial supernatant was collected. The nuclear fraction was further processed and washed twice with a hypotonic buffer (20 mM Tris-HCl pH 7.4, 3 mM MgCl_2_, and 10 mM NaCl) followed by centrifugation at 1,000 *g*. The pellet was then resuspended in nuclear extraction buffer (20 mM Tris-HCl pH 7.9, 400 mM NaCl, 1 mM EDTA, 1 mM EGTA, and 1% Nonidet P-40) and incubated on ice for 45 min with gentle vortexing every 5 min. Following incubation, the lysate was centrifuged at 14,000 *g* to remove debris, and the nuclear fraction was collected from the supernatant. Protein estimation in all cellular fractions was conducted using the BCA method. The purity of the fractions was confirmed by immunoblotting with Lamin B1 (nuclear marker), ATP5A (mitochondrial marker), and GAPDH (cytosolic/post-mitochondrial marker).

#### For liver tissue

Liver tissue was finely minced and resuspended in resuspension buffer (250 mM sucrose, 250 mM mannitol, 25 mM HEPES pH 7.4, 10 mM KCl, 0.25 mM EDTA, 10 mM EGTA, 1.5 mM MgCl₂, 1 mM DTT, 0.1% BSA, and 1X protease inhibitor cocktail). The tissue suspension was homogenized using a Dounce homogenizer with 20 strokes of Pestle A followed by 20 strokes of Pestle B (repeated twice). The homogenate was first centrifuged at 2,500 *g* for 10 min at 4°C to obtain the nuclear fraction. The post-nuclear supernatant was subsequently centrifuged at 7,000 *g* for 15 min at 4°C to isolate the mitochondrial pellet, which was washed and resuspended in the same buffer, without chelating agents, at a final protein concentration of 1 mg/mL. The nuclear fraction was resuspended in hypotonic buffer (20 mM Tris-HCl pH 7.4, 3 mM MgCl_2_, and 10 mM NaCl), and centrifuged at 300 *g* twice to remove debris. The resulting supernatant was further centrifuged at 700 *g* to isolate intact nuclei. The nuclear pellet was resuspended in hypotonic buffer (containing 1% Nonidet P-40) and incubated on ice for 15 min with vortexing every 5 min. Following incubation, the sample was centrifuged at 700 *g*, and the pellet was washed with the hypotonic buffer. The washed nuclear pellet was then resuspended in hypertonic lysis buffer (20 mM Tris-HCl pH 7.9, 400 mM NaCl, 1 mM EDTA, 1 mM EGTA, and 1% Nonidet P-40), and incubated on ice for 45 min with gentle vortexing every 5 min. After incubation, the lysate was centrifuged at 14,000 *g* to pellet chromatin, and the supernatant containing the soluble nuclear fraction was collected for further analyses. Estimation of protein concentration and purity of the fractions was conducted as described for cell line fractionation.

#### PolyP extraction from mammalian samples

PolyP from the nuclear fraction, mitochondrial fraction, and post-mitochondrial supernatant was extracted using the phenol-chloroform method. Each fraction was treated with TEELS buffer (10 mM Tris-Cl pH 8.0, 10 mM EDTA, 10 mM EGTA, 100 mM LiCl, 0.2% SDS, and 5 mM NaF) and an equal volume of acid phenol (pH 4.8). Samples were vortexed for 3 min and centrifuged at 18,000 *g* for 10 min at 25°C. Following centrifugation, the upper aqueous phase was carefully collected without disturbing the phenol layer or the protein ring at the interface. The aqueous phase was then washed with an excess volume of chloroform and centrifuged again at 18,000 *g* for 15 min. The resulting aqueous layer was collected, mixed with 2.5 volumes of 100% ethanol, and incubated at –80 °C overnight to precipitate polyP. The next day, samples were centrifuged at 18,000 *g* for 30 min at 4°C. The resulting white or translucent pellet containing polyP was resuspended in 200 µL of recording buffer (150 mM KCl, and 20 mM HEPES, pH 7.0).

### Treatment of cells prior to polyP extraction

U-2 OS cells were seeded and allowed to adhere for 24 h prior to treatment. To alter glucose in the growth media, cells were shifted to DMEM without glucose (Himedia, AL186) supplemented with 25 mM glucose (high glucose), 5 mM glucose (low glucose), or 10 mM galactose for 9 h prior to harvest. For treatment with inhibitors, cells seeded in normal growth medium were shifted to low-glucose medium for 9 h. Following this pre-incubation, cells were treated with either Oligomycin (100 ng/mL; Sigma-Aldrich, 75351), Antimycin A (2 µM; Sigma-Aldrich, A8674), FCCP (2 µM; Sigma-Aldrich, C2920), or DMSO (vehicle control) for an additional 3 h in the same low-glucose medium. After the treatment, cells were harvested for mitochondrial polyP extraction and quantification. To inhibit IP6K activity, U-2 OS cells were seeded and allowed to adhere for 24 h prior to treatment. The medium was changed to low-glucose DMEM and cells were treated with 10 µM TNP (Sigma-Aldrich, T3955) or DMSO (vehicle control) for 12 h. After the treatment, cells were harvested for mitochondrial polyP extraction and quantification.

### Quantification of polyP_20_ by malachite green assay

Hexahistidine-tagged *Saccharomyces cerevisiae* exopolyphosphatase (*Sc*PPX) used for the biochemical quantification of polyP (encoded on plasmid pTrcHisB) was expressed in *E. coli* (BL21DE3), and purified by affinity chromatography using a Ni-NTA HiTrap column on an FPLC system (Cytiva Akta pure). The length of polyP in sodium hexametaphosphate (Cat. No. P8510 Sigma Aldrich) was estimated to be an average chain length of ∼20 Pi residues, by comparison with short- (polyP_14_), medium- (polyP_60_) and long- (polyP_130_) chain polyP from RegeneTiss (Cat. Nos. 638-51691), resolved on a 15% polyacrylamide gel (Fig. S1A). We used this polyP_20_ as a standard for all polyP quantification assays. The linear nature of polyP_20_ chains was confirmed by digestion with *Sc*PPX (Fig. S1A). The amount of polyP_20_ in Pi terms was quantified using the malachite green method [58]. 100 mg polyP_20_ was dissolved in 1 mL nuclease free water, and further diluted to 500 µg/mL and 750 µg/mL in recording buffer. 50 µL of these solutions were treated with 2 µg of *Sc*PPX for 2 h at 37°C. To prepare the reagent mix for the malachite green assay, Solution A (0.45% malachite green (Sigma Aldrich Cat. No. 213020) in distilled water), and Solution B (4.2% ammonium molybdate tetrahydrate (Sigma Aldrich Cat No. A7302) in 4N HCl) stored at 4°C were mixed in a 3:1 ratio and filtered through Whatmann paper grade 1. K_2_HPO_4_ (ranging from 200 pmole to 1 nmole) was used as a standard. The standard and samples in a volume of up to 10 µL were added with 200 µL reagent mix, incubated in the dark for 5 min with shaking, and the absorbance was measured at 660 nm. The amount of Pi released from polyP_20_ samples treated with *Sc*PPX was estimated from the linear regression standard curve, and used to determine the concentration of the polyP_20_ stock in Pi terms.

### Quantification of polyP using DAPI

PolyP_20_ (ranging from 300 to 10,000 pmoles in Pi equivalents) was used as a standard for quantification of polyP. For biological samples, the extract was divided into two equal parts - one part was treated with 1 µg of *Sc*PPX, while the other remained untreated. Both aliquots were incubated at 37°C for 16 h. Standards and samples in a volume of 100 µL recording buffer (150 mM KCl, and 20 mM HEPES, pH 7.0) were mixed with an equal volume of 60 µM DAPI, and were incubated at room temperature in the dark for 15 min. Fluorescence was recorded using a multimode plate reader (Perkin Elmer EnSpire; Ex 415 nm, Em 550 nm). The polyP_20_ equivalents in the untreated and *Sc*PPX-treated aliquots were interpolated from the linear regression standard curve, and the difference in the interpolated values corresponded to the amount of polyP present in the sample.

### *Ex-vivo* polyP synthesis assay

Mitochondria were isolated from liver tissue of *Ip6k1*⁺^/^⁺ and *Ip6k1*⁻^/^⁻ mice, or from U-2 OS cell lines as described above, and resuspended in KCl buffer (10 mM Tris-Cl pH 7.1, 120 mM KCl, and 1 mM EGTA). Protein content in the sample was estimated using the Pierce™ Bradford Protein Assay Kit (Thermo Scientific, 1856210). The integrity of the mitochondrial fraction was assessed using JC-1 dye [59]. For JC-1 fluorescence based detection of membrane potential of isolated mitochondria, 5 µg of mitochondria were incubated in 200 µL KCl buffer containing 5 mM glutamate, 5 mM malate, 10 mM succinate, and 0.5 µM JC-1 dye. PerkinElmer EnSpire multimode plate reader was used to detect the JC-1 dye fluorescence emission spectrum from 500 to 640 nm at an interval of 10 nm (Ex 488 nm). The polyP synthesis assay was conducted in a final volume of 200 µL KCl buffer, and contained isolated mitochondria (0.5 mg/mL), 30 µM DAPI, 5 mM glutamate, 5 mM malate, 10 mM succinate, and the indicated concentrations of ATP and K_2_HPO_4_ in a 96 well black plate (Thermo Scientific, Cat no. 237108). DAPI fluorescence was recorded at 1 min intervals (Ex 415 nm, Em 550 nm). The relative DAPI fluorescence over time was calculated for each well as F_t_/F_0_ (where F_t_ is the fluorescence at time t and F_0_ is fluorescence at time zero). Where indicated, AMP-PNP, antimycin A, FCCP, or oligomycin were added during the polyP synthesis assay.

### Analysis of cellular inositol pyrophosphates

NTC, *IP6K1*^-/-^ and *IP6K1*^-/-^ U-2 OS cells stably expressing active or catalytically inactive IP6K1 (K226A/S334A) were labeled with [^3^H]-inositol as described earlier [54, 60]. Cells seeded in 60 mm dishes in normal growth medium were allowed to attain 30% confluence and then transferred to inositol-free DMEM (MP Biomedicals, D9802-06.25) containing 10% dialysed FBS, and labeled with 40 μCi *myo*-2-[^3^H] inositol for 2.5 days. The media was removed and fresh media containing *myo*-2-[^3^H] inositol (40 μCi) was added for another 2.5 days. At the end of 5 days, when isotopic labeling was achieved, cells were washed and collected by scraping in chilled PBS. From the cell pellet, soluble inositol phosphates were extracted by the addition of 350 μL extraction buffer (0.6 M HClO_4_, 2 mM EDTA, and 0.2 mg/mL phytic acid) on ice for 15-20 min, followed by centrifugation at 21,000 *g* for 10 min. The supernatant containing soluble inositol phosphates was collected, and lipid inositols in the pellet were extracted with 1 mL lipid extraction buffer (0.1 N NaOH and 0.1% Triton X-100) at room temperature with end-over-end mixing for 4-5 h. The soluble inositol phosphate extract was mixed with ∼120 μL neutralization solution (1 M K_2_CO_3_ and 5 mM EDTA). Tubes were left open on ice for 1 h, followed by centrifugation at 21,000 *g* for 10 min at 4°C. The extracted inositol phosphates were resolved by HPLC (5125 HPLC pumps, Waters) on a Partisphere SAX column (4.6 mm × 125 mm, HiChrome) using a gradient of Buffer A (1mM EDTA) and Buffer B (1mM EDTA and 1.3 M (NH_4_)_2_HPO_4_; pH 3.8) as follows: 0-5 min, 0% B; 5-10 min, 0-20% B; 10-70 min, 20-100% B; 70-80 min, 100% B. 1 mL fractions containing soluble inositol phosphates were mixed with 3 mL scintillation cocktail (Ultima-Flo AP) and counted for 5 min in a liquid scintillation counter (Tri-Carb 2910 TR, Perkin Elmer).

### Immunofluorescence analysis

Cells grown on glass coverslips in a 24-well plate were incubated with 300 nM Mitotracker Red (Invitrogen, M22425) for 1 h in DMEM without FBS. Post-incubation, cells were washed and incubated with pre-warmed DMEM with 10% FBS for 30 min to remove excess Mitotracker Red. Cells were subsequently washed with PBS and fixed with 4% formaldehyde for 10 min at room temperature, and permeabilised with 0.15% Triton X-100 for 10 min at room temperature. Non-specific antibody binding was blocked by incubating the cells in blocking buffer (3% BSA in PBS with 0.15% Triton X-100) for 1 h at room temperature. Cells were incubated overnight at 4°C with an anti-IP6K1 antibody (Sigma-Aldrich, HPA040825; 1:300) diluted in blocking buffer. The next day, cells were washed three times with PBS and incubated with a fluorophore-conjugated secondary antibody (Alexa Fluor 488 anti-rabbit IgG; 1:500) diluted in blocking buffer for 1 h at room temperature. Cells were washed three times with PBS and the coverslips were mounted on glass slides using an antifade mounting medium containing DAPI (H-1200, Vecta Labs), air dried, and sealed. Images were acquired using a Zeiss LSM 700 confocal microscope equipped with 405, 488, and 555 nm lasers and, fitted with a 63×1.4 NA oil-immersion objective. Images are presented as maximum intensity projections (MIP) of z-stacks in the xy plane, processed using ZEN software. Pearson correlation coefficient analysis was performed using ZEN Black software.

### Analysis of mitochondrial respiration using Seahorse assay

Mitochondrial respiration (oxygen consumption rate; OCR) in U-2 OS cells was analyzed using the Seahorse extracellular flux analyzer (Agilent Technologies) [47, 61]. Briefly, 10,000 cells were seeded per well in a Seahorse 24-well assay plate in growth medium (DMEM, 10% FBS, 2 mM L-glutamine) supplemented with 25 mM glucose. After 24 h, cells were switched to growth medium containing high glucose (25 mM) or low glucose (5 mM) for a further 12 h. Cells were washed twice with 1 mL pre-warmed OCR XF base medium (pH 7.4) (Agilent Technologies, Cat. No 00840-000) supplemented with 2 mM L-glutamine, 1 mM sodium pyruvate, and either 25 mM (high) or 5 mM (low) glucose, and equilibriated in 500 µL of the same medium for 45 min at 37°C in a non-CO_2_ incubator. OCR was recorded under basal conditions, and following sequential injections of 2 µM oligomycin (ATP synthase inhibitor), 2 µM FCCP (mitochondrial uncoupler), and a combination of 0.05 µM rotenone (complex I inhibitor) and 4 µM antimycin A (complex III inhibitor). OCR data were normalized to the protein content of each well, as quantified using BCA kit (Thermo fisher scientific Cat no A55860) at the end of the assay.

### Detection of mitochondrial membrane potential using TMRM-based flow cytometry

To assess mitochondrial membrane potential (ΔΨm), U-2 OS cells were cultured in DMEM supplemented with 10% FBS. 24 h post-seeding, cells were incubated with 300 nM TMRM for 30 min at 37°C in the dark. Cells were washed with PBS, trypsinized, and resuspended in DMEM containing 10% FBS to neutralize trypsin activity. The cells were further centrifuged, washed, and resuspended in PBS. Subsequently, flow cytometry analysis of TMRM-stained cells was performed using a 488 nm excitation laser and a 585/42 nm emission filter on a BD LSRFortessa X-20 flow cytometer. Mitochondria with intact membrane potential exhibit higher TMRM fluorescence intensity, whereas depolarized mitochondria show reduced fluorescence.

### Statistical analysis

GraphPad Prism 8 was used to perform statistical analyses and prepare graphs. The number of biologically independent replicates (N) for each experiment are indicated in the figure legends. P-values are from a one-sample t-test or Student’s t-test as indicated and P≤0.05 was considered statistically significant.

## Supporting information

PolyP8 mass Spectrometry data

CRAPome analysis

DAVID functional annotation clustering

## AUTHOR CONTRIBUTIONS

**Conceptualization**: J.S.L., U.K., M. J., and R.B.; **Methodology**: J.S.L. and R.B.; **Validation**: J.S.L. and A.K.; **Formal analysis**: J.S.L. and A.K.; **Investigation:** J.S.L., A.K., and A.S.**; Resources:** J.S., N.S., H.J.J.: **Writing - original draft**: J.S.L., A.K., and R.B.; **Writing - review and editing**: J.S.L. and R.B.; **Visualization**: J.S.L.; **Supervision**: R.B; **Project administration**: R.B.; **Funding acquisition**: H.J.J., M.J., R.B. All authors have read and agreed to the published version of the manuscript.

## ACKNOWLEDGEMENTS

We thank Manasa Chanduri and Debaditya De for polyP pull-down and mass spectrometry data; Subhash Khatri for Seahorse assays; Manasa Chanduri for DAPI emission spectra; Akruti Shah for the generation of U-2 OS NTC and *IP6K1*^-/-^ cells; Shubhra Ganguli for tritium inositol labeling; the staff at the Sophisticated Equipment Facility and Experimental Animal Facility at CDFD for technical assistance. We also thank Swasti Raychaudhuri, R. Harinarayanan, Naresh Sepuri, and the members of the Laboratory of Cell Signalling, CDFD for their valuable feedback.

## DATA AVAILABILITY STATEMENT

The mass spectrometry data used in this manuscript have been deposited to the MassIVE, a full member of the Proteome Xchange Consortium, and can be accessed through Data set identifier ID-MSV000098211 (https://massive.ucsd.edu).

---

**Figure S1.**
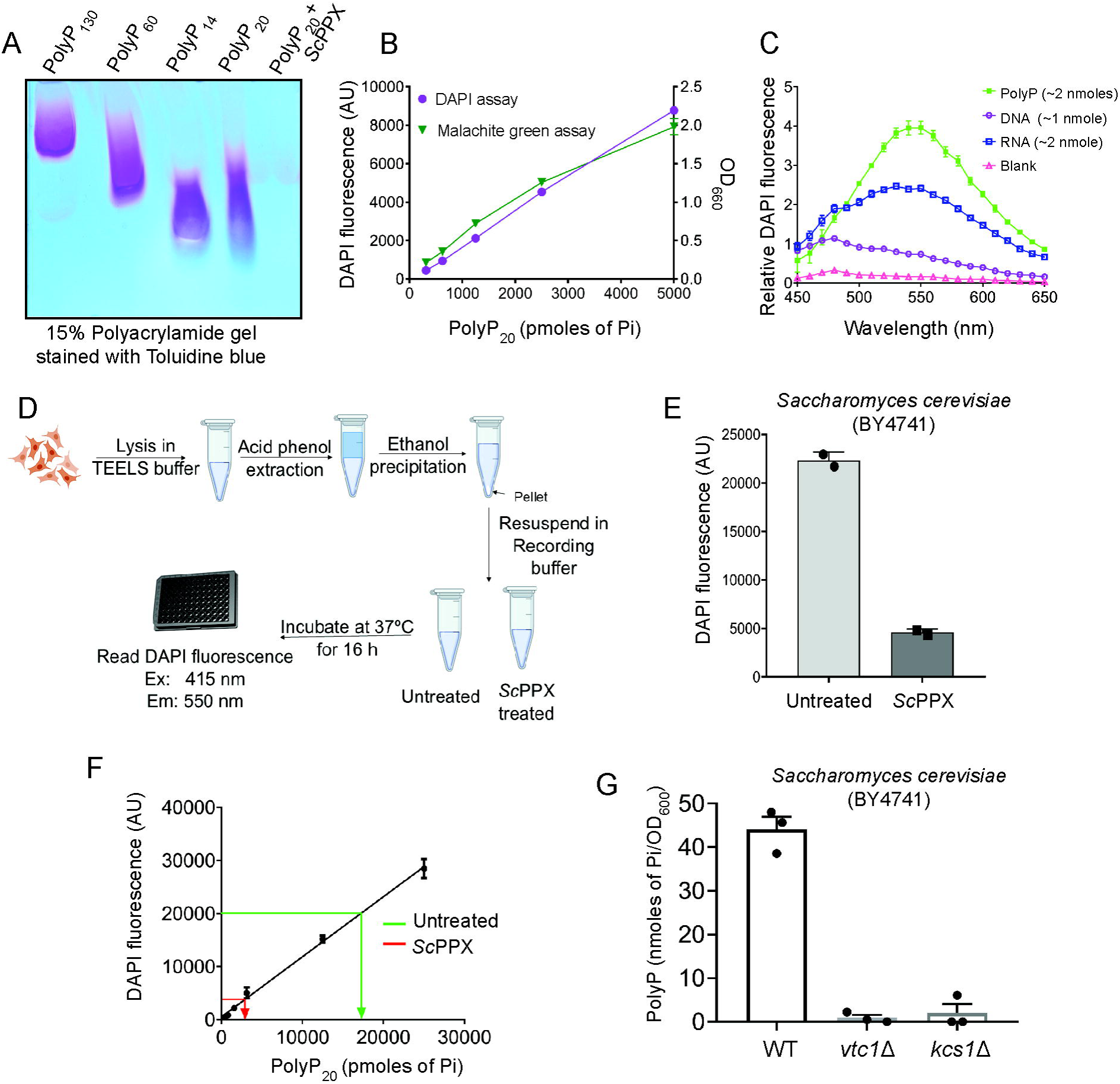
PolyP quantification by DAPI. **(A)** 15% Tris-borate-EDTA (TBE) polyacrylamide gel stained with toluidine blue representing polyP standards (polyP_130_, polyP_60_, and polyP_14_) from RegeneTiss along with untreated or *Sc*PPX-treated sodium hexametaphosphate, estimating the chain length of the latter to be approximately 25 Pi units (polyP_20_), and demonstrating its linear nature. **(B)** Standard curve for polyP_20_ quantification using Malachite green assay (right Y- axis) and DAPI fluorescence (left Y-axis). Malachite green absorbance (OD: 660 nm) and DAPI fluorescence intensity (Ex: 415 nm, Em: 550 nm) were measured in the presence of increasing amount of polyP_20_ (range: 156 pmoles to 5000 pmoles of Pi). **(C)** Fluorescence emission spectra of DAPI fluorescence obtained upon excitation at 415 nm, with emission measured between 450 nm and 650 nm. Distinct emission profiles were observed for DAPI bound to DNA (purple), RNA (blue), and polyP_20_ (green), compared with blank (pink). Emission maximum was observed at 480 nm for DAPI-DNA and 550 nm for DAPI-RNA and DAPI-polyP_20_. **(D)** Schematic of protocol for polyP extraction and purification, followed by *Sc*PPX treatment and detection by DAPI. **(E)** Quantification of polyP extracted from *S. cerevisiae* (strain BY4741). Bar graph represents DAPI fluorescence intensity for untreated and *Sc*PPX-treated polyP extracts (Ex 415 nm, Em 550 nm). **(F)** Standard curve for polyP_20_ with relative DAPI fluorescence intensities representing the interpolation of the DAPI fluorescence values from bar graph in (E). **(G)** Bar graph representing polyP levels in WT, *vtc1*Δ and *kcs1*Δ *S. cerevisiae* (strain BY4741) normalised to 1 OD culture.

**Figure S2.**
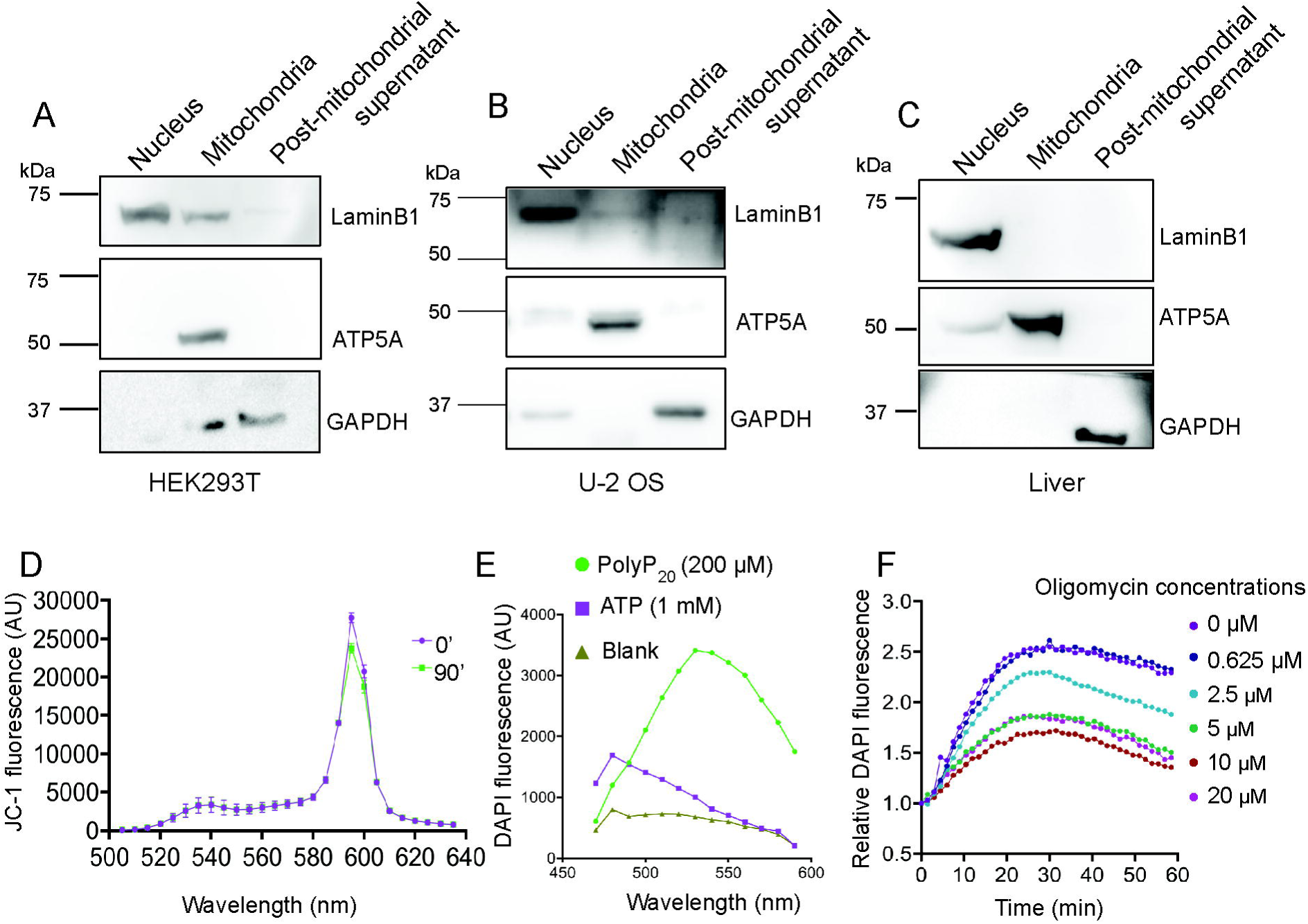
**(A-C)** Representative immunoblots examining subcellular fractionation of HEK293T (A), U-2 OS cells (B) and mouse liver (C). The nuclear, mitochondrial, and post-mitochondrial fractions were marked by the enrichment of Lamin B1, ATP5A, and GAPDH, respectively. **(D)** JC-1 dye fluorescence spectra of mitochondria isolated from mouse liver incubated in KCl buffer containing glutamate (5 mM), malate (5 mM), and succinate (5 mM), with 0.5 μM JC-1 dye. Spectra were recorded at 0 min (magenta) and 90 min (green) post-incubation. Samples were excited at 488 nm, and emission was recorded from 500 to 640 nm in 10 nm intervals. **(E)** DAPI emission spectra for polyP (200 μM) and ATP (1 mM) upon excitation at 415 nm. **(F)** Line graph depicting the fold change in DAPI fluorescence as a measure of polyP synthesis in isolated mitochondria incubated with glutamate (5 mM), malate (5 mM), succinate (10 mM), Pi (5 mM), and ATP (1 mM) under varying concentrations of oligomycin.

**Figure S3.**
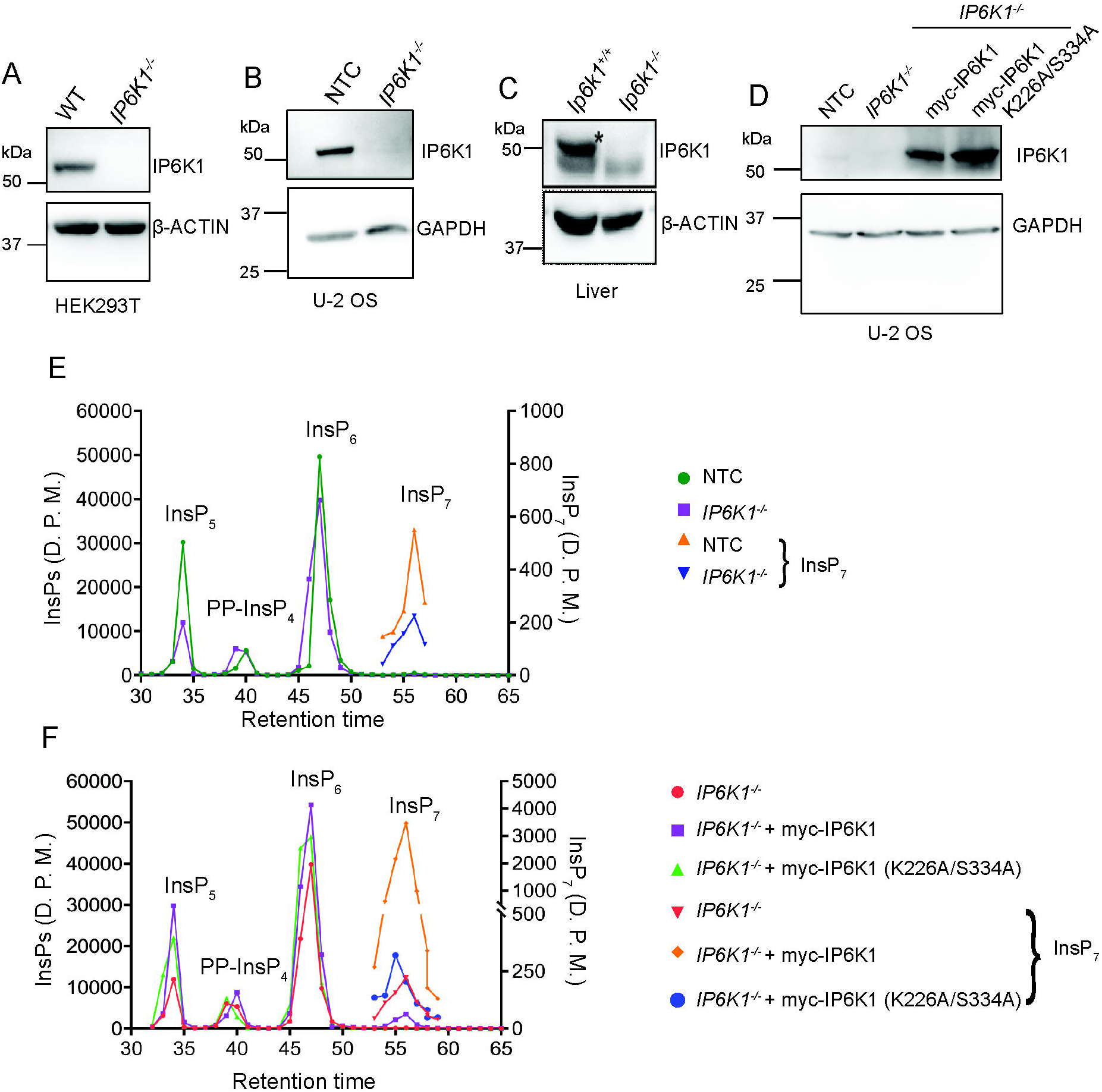
Characterisation of IP6K1 knock out cell lines and mouse liver. (A-C) Representative immunoblots showing the absence of IP6K1 in *IP6K1^-/-^* knockout HEK293T (A), and U-2 OS (B) cell lines, and *Ip6k1^-/-^* mouse liver compared with their respective controls (C). **(D)** Representative immunoblot demonstrating the stable expression of myc-tagged active or catalytically inactive (K226A/S334A) mouse IP6K1 in *IP6K1^-/-^* U-2 OS cell line. **(E-F)** HPLC profiles of [^3^H]-inositol-labelled NTC and *IP6K1^-/-^* U-2 OS cells (E), and *IP6K1^-/-^* U-2 OS cells stably expressing myc-tagged active or catalytically inactive (K226A/S334A) mouse IP6K1 (F). The profile for InsP_7_ is shown separately (right Y axis), in addition to all InsPs (left Y axis).

**Figure S4.**
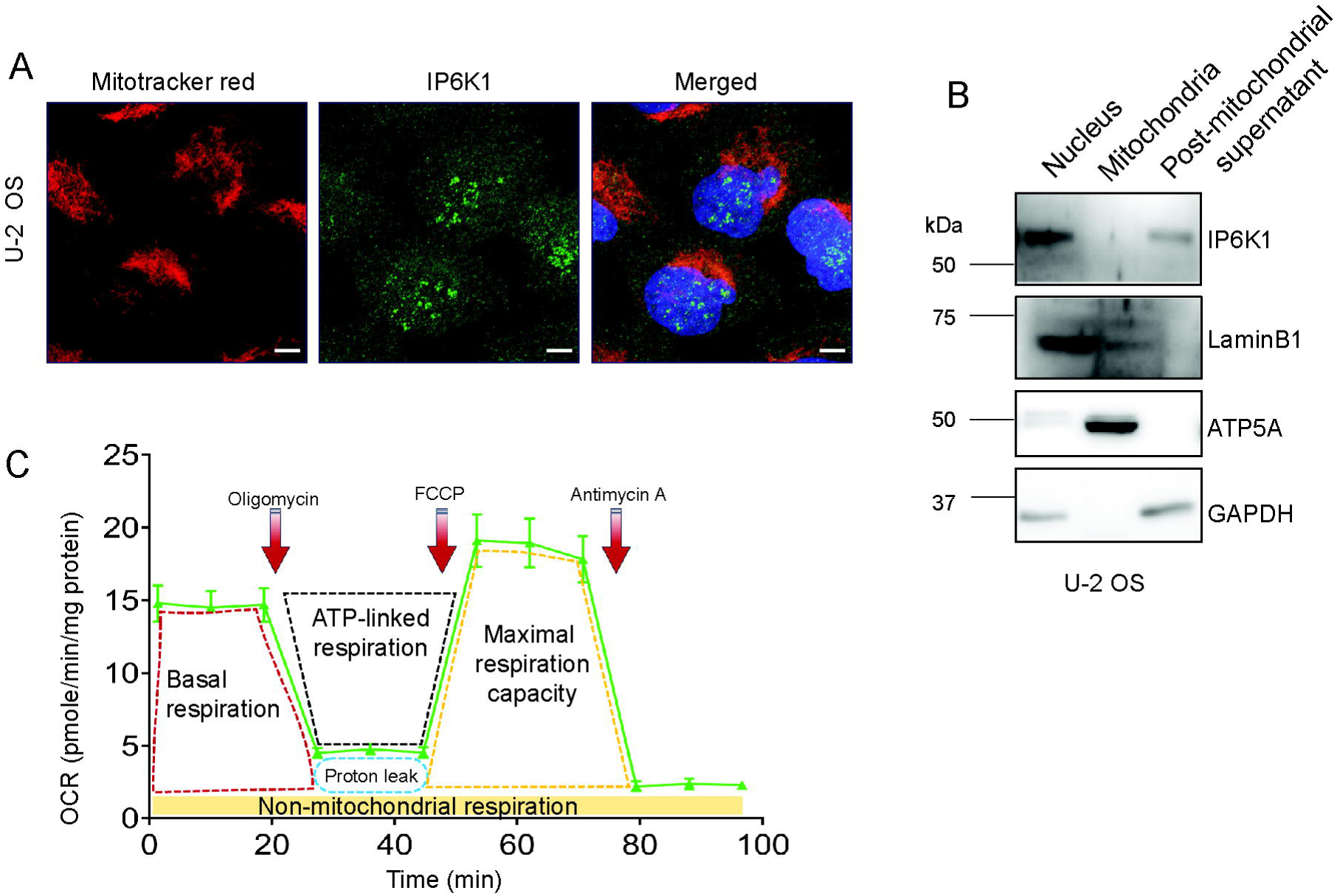
IP6K1 does not localise to mitochondria. **(A)** Asynchronous U-2 OS cells were stained with Mitotracker Red (red) and anti-IP6K1 antibody (green): scale bars, 5 µm. Co-localization of Mitotracker Red with IP6K1 was assessed by calculating Pearsons’s correlation coefficient, which yielded a value of 0.069 ± 0.056 (mean ± S.E.M.), indicating no significant co-localisation. **(B)** Representative immunoblot showing the enrichment of IP6K1 in the nuclear and post-mitochondrial supernatant fractions, with no detectable signal in the mitochondrial fraction. **(C)** Schematic representation of interpretation of OCR traces, expressed as pmoles O_2_/min/µg protein. Arrows indicate the time points at which Oligomycin, FCCP, and Antimycin A/Rotenone are added. Baseline cellular OCR is measured initially, from which basal respiration is calculated by subtracting non-mitochondrial respiration. Upon addition of oligomycin (Complex V inhibitor), the decrease in OCR is used to determine ATP-linked respiration (by subtracting the oligomycin-inhibited rate from the baseline OCR). Proton leak respiration is calculated by subtracting non-mitochondrial respiration from the oligomycin-inhibited OCR. Next, FCCP (a protonophore), is added to collapse the inner mitochondrial membrane gradient, allowing the ETC to function at its maximal rate. Maximal respiratory capacity is calculated by subtracting non-mitochondrial respiration from the FCCP-stimulated OCR. Lastly, Antimycin A and Rotenone, inhibitors of complex III and I respectively, are added to shut down ETC function, revealing the non-mitochondrial respiration.

## REFERENCES

1. Albi T. and Serrano A. (2016) Inorganic polyphosphate in the microbial world. Emerging roles for a multifaceted biopolymer. World J Microbiol Biotechnol 32, 27.

2. Borghi F. and Saiardi A. (2023) Evolutionary perspective on mammalian inorganic polyphosphate (polyP) biology. Biochem Soc Trans 51, 1947–56.

3. Khan A., Mallick M., Ladke J.S. and Bhandari R. (2024) The ring rules the chain - inositol pyrophosphates and the regulation of inorganic polyphosphate. Biochem Soc Trans 52, 567–80.

4. Kumble K.D. and Kornberg A. (1995) Inorganic polyphosphate in mammalian cells and tissues. J Biol Chem 270, 5818–22.

5. Pisoni R.L. and Lindley E.R. (1992) Incorporation of [32P]orthophosphate into long chains of inorganic polyphosphate within lysosomes of human fibroblasts. J Biol Chem 267, 3626–31.

6. Jimenez-Nunez M.D., Moreno-Sanchez D., Hernandez-Ruiz L., Benitez-Rondan A., Ramos-Amaya A., Rodriguez-Bayona B., et al. (2012) Myeloma cells contain high levels of inorganic polyphosphate which is associated with nucleolar transcription. Haematologica 97, 1264–71.

7. Ruiz F.A., Lea C.R., Oldfield E. and Docampo R. (2004) Human platelet dense granules contain polyphosphate and are similar to acidocalcisomes of bacteria and unicellular eukaryotes. J Biol Chem 279, 44250–7.

8. Smith S.A., Mutch N.J., Baskar D., Rohloff P., Docampo R. and Morrissey J.H. (2006) Polyphosphate modulates blood coagulation and fibrinolysis. Proc Natl Acad Sci U S A 103, 903–8.

9. Seidlmayer L.K., Gomez-Garcia M.R., Blatter L.A., Pavlov E. and Dedkova E.N. (2012) Inorganic polyphosphate is a potent activator of the mitochondrial permeability transition pore in cardiac myocytes. J Gen Physiol 139, 321–31.

10. Solesio M.E., Garcia Del Molino L.C., Elustondo P.A., Diao C., Chang J.C. and Pavlov E.V. (2020) Inorganic polyphosphate is required for sustained free mitochondrial calcium elevation, following calcium uptake. Cell Calcium 86, 102127.

11. Wang Y., Li M., Li P., Teng H., Fan D., Du W., et al. (2019) Progress and applications of polyphosphate in bone and cartilage regeneration. Biomed Res Int 2019, 5141204.

12. Abramov A.Y., Fraley C., Diao C.T., Winkfein R., Colicos M.A., Duchen M.R., et al. (2007) Targeted polyphosphatase expression alters mitochondrial metabolism and inhibits calcium-dependent cell death. Proc Natl Acad Sci U S A 104, 18091–6.

13. Ahn K. and Kornberg A. (1990) Polyphosphate kinase from *Escherichia coli.* Purification and demonstration of a phosphoenzyme intermediate. J Biol Chem 265, 11734–9.

14. Hothorn M., Neumann H., Lenherr E.D., Wehner M., Rybin V., Hassa P.O., et al. (2009) Catalytic Core of aMembrane-Associated Eukaryotic Polyphosphate Polymerase. Science 324, 513–5.

15. Lynn W.S. and Brown R.H. (1963) Synthesis of polyphosphate by rat liver mitochondria. Biochem Biophys Res Commun 11, 367–71.

16. Pavlov E., Aschar-Sobbi R., Campanella M., Turner R.J., Gomez-Garcia M.R. and Abramov A.Y. (2010) Inorganic polyphosphate and energy metabolism in mammalian cells. J Biol Chem 285, 9420–8.

17. Baev A.Y., Angelova P.R. and Abramov A.Y. (2020) Inorganic polyphosphate is produced and hydrolyzed in F0F1-ATP synthase of mammalian mitochondria. Biochem J 477, 1515–24.

18. Auesukaree C., Tochio H., Shirakawa M., Kaneko Y. and Harashima S. (2005) Plc1p, Arg82p, and Kcs1p, enzymes involved in inositol pyrophosphate synthesis, are essential for phosphate regulation and polyphosphate accumulation in Saccharomyces cerevisiae. J Biol Chem 280, 25127–33.

19. Gerasimaite R., Pavlovic I., Capolicchio S., Hofer A., Schmidt A., Jessen H.J., et al. (2017) Inositol Pyrophosphate Specificity of the SPX-Dependent Polyphosphate Polymerase VTC. ACS Chem Biol 12, 648–53.

20. Wild R., Gerasimaite R., Jung J.Y., Truffault V., Pavlovic I., Schmidt A., et al. (2016) Control of eukaryotic phosphate homeostasis by inositol polyphosphate sensor domains. Science 352, 986–90.

21. Ghosh S., Shukla D., Suman K., Lakshmi B.J., Manorama R., Kumar S., et al. (2013) Inositol hexakisphosphate kinase 1 maintains hemostasis in mice by regulating platelet polyphosphate levels. Blood 122, 1478–86.

22. Hou Q., Liu F., Chakraborty A., Jia Y., Prasad A., Yu H., et al. (2018) Inhibition of IP6K1 suppresses neutrophil-mediated pulmonary damage in bacterial pneumonia. Sci Transl Med 10.

23. Bentley-DeSousa A., Holinier C., Moteshareie H., Tseng Y.C., Kajjo S., Nwosu C., et al. (2018) A Screen for Candidate Targets of Lysine Polyphosphorylation Uncovers a Conserved Network Implicated in Ribosome Biogenesis. Cell Rep 22, 3427–39.

24. Azevedo C., Singh J., Steck N., Hofer A., Ruiz F.A., Singh T., et al. (2018) Screening a Protein Array with Synthetic Biotinylated Inorganic Polyphosphate To Define the Human PolyP-ome. ACS Chem Biol 13, 1958–63.

25. Neville N., Lehotsky K., Yang Z., Klupt K.A., Denoncourt A., Downey M., et al. (2023) Modification of histidine repeat proteins by inorganic polyphosphate. Cell Rep 42, 113082.

26. Singh J., Steck N., De D., Hofer A., Ripp A., Captain I., et al. (2019) A Phosphoramidite Analogue of Cyclotriphosphate Enables Iterative Polyphosphorylations. Angew Chem Int Ed Engl 58, 3928–33.

27. Mellacheruvu D., Wright Z., Couzens A.L., Lambert J.P., St-Denis N.A., Li T., et al. (2013) The CRAPome: a contaminant repository for affinity purification-mass spectrometry data. Nat Methods 10, 730–6.

28. Huang D.W., Sherman B.T. and Lempicki R.A. (2009) Systematic and integrative analysis of large gene lists using DAVID bioinformatics resources. Nature Protocols 4, 44–57.

29. Gomez Garcia M.R. (2014) Extraction and Quantification of Poly P, Poly P Analysis by Urea-PAGE. Bio Protoc 4.

30. Lazaro B., Sarrias A., Tadeo F.J., Marc Martinez-Lainez J., Fernandez A., Quandt E., et al. (2025) Optimized biochemical method for human Polyphosphate quantification. Methods 234, 211–22.

31. Neef D.W. and Kladde M.P. (2003) Polyphosphate loss promotes SNF/SWI- and Gcn5-dependent mitotic induction of PHO5. Mol Cell Biol 23, 3788–97.

32. Bru S., Jimenez J., Canadell D., Arino J. and Clotet J. (2016) Improvement of biochemical methods of polyP quantification. Microb Cell 4, 6–15.

33. Bondy-Chorney E., Abramchuk I., Nasser R., Holinier C., Denoncourt A., Baijal K., et al. (2020) A Broad Response to Intracellular Long-Chain Polyphosphate in Human Cells. Cell Rep 33, 108318.

34. Aschar-Sobbi R., Abramov A.Y., Diao C., Kargacin M.E., Kargacin G.J., French R.J., et al. (2008) High sensitivity, quantitative measurements of polyphosphate using a new DAPI-based approach. J Fluoresc 18, 859–66.

35. Martin P. and Van Mooy B.A. (2013) Fluorometric quantification of polyphosphate in environmental plankton samples: extraction protocols, matrix effects, and nucleic acid interference. Appl Environ Microbiol 79, 273–81.

36. Werner T.P., Amrhein N. and Freimoser F.M. (2005) Novel method for the quantification of inorganic polyphosphate (iPoP) in Saccharomyces cerevisiae shows dependence of iPoP content on the growth phase. Arch Microbiol 184, 129–36.

37. Solesio M.E., Elustondo P.A., Zakharian E. and Pavlov E.V. (2016) Inorganic polyphosphate (polyP) as an activator and structural component of the mitochondrial permeability transition pore. Biochem Soc Trans 44, 7–12.

38. Liberti M.V. and Locasale J.W. (2016) The Warburg Effect: How Does it Benefit Cancer Cells? Trends Biochem Sci 41, 211–8.

39. Staricha K., Meyers N., Garvin J., Liu Q., Rarick K., Harder D., et al. (2020) Effect of high glucose condition on glucose metabolism in primary astrocytes. Brain Res 1732, 146702.

40. Shiratori R., Furuichi K., Yamaguchi M., Miyazaki N., Aoki H., Chibana H., et al. (2019) Glycolytic suppression dramatically changes the intracellular metabolic profile of multiple cancer cell lines in a mitochondrial metabolism-dependent manner. Sci Rep 9, 18699.

41. Aguer C., Gambarotta D., Mailloux R.J., Moffat C., Dent R., McPherson R., et al. (2011) Galactose enhances oxidative metabolism and reveals mitochondrial dysfunction in human primary muscle cells. PLoS One 6, e28536.

42. Taira Y., Okegawa Y., Sugimoto K., Abe M., Miyoshi H. and Shikanai T. (2013) Antimycin A-like molecules inhibit cyclic electron transport around photosystem I in ruptured chloroplasts. FEBS Open Bio 3, 406–10.

43. Wang S.H., Tung T.H., Chiu S.P., Chou H.Y., Hung Y.H., Lai Y.T., et al. (2021) Detecting Effects of Low Levels of FCCP on Stem Cell Micromotion and Wound-Healing Migration by Time-Series Capacitance Measurement. Sensors (Basel) 21.

44. Symersky J., Osowski D., Walters D.E. and Mueller D.M. (2012) Oligomycin frames a common drug-binding site in the ATP synthase. Proc Natl Acad Sci U S A 109, 13961–5.

45. Shah A. and Bhandari R. (2021) IP6K1 upregulates the formation of processing bodies by influencing protein-protein interactions on the mRNA cap. J Cell Sci 134.

46. Szijgyarto Z., Garedew A., Azevedo C. and Saiardi A. (2011) Influence of inositol pyrophosphates on cellular energy dynamics. Science 334, 802–5.

47. Brand M.D. and Nicholls D.G. (2011) Assessing mitochondrial dysfunction in cells. Biochem J 435, 297–312.

48. Ganguli S., Shah A., Hamid A., Singh A., Palakurti R. and Bhandari R. (2020) A high energy phosphate jump - From pyrophospho-inositol to pyrophospho-serine. Adv Biol Regul 75, 100662.

49. Shears S.B. (2009) Diphosphoinositol polyphosphates: metabolic messengers? Mol Pharmacol 76, 236–52.

50. Wundenberg T. and Mayr G.W. (2012) Synthesis and biological actions of diphosphoinositol phosphates (inositol pyrophosphates), regulators of cell homeostasis. Biol Chem 393, 979–98.

51. Rajasekaran S.S., Kim J., Gaboardi G.C., Gromada J., Shears S.B., Dos Santos K.T., et al. (2018) Inositol hexakisphosphate kinase 1 is a metabolic sensor in pancreatic beta-cells. Cell Signal 46, 120–8.

52. Morgan J.A.M., Singh A., Kurz L., Nadler-Holly M., Ruwolt M., Ganguli S., et al. (2024) Extensive protein pyrophosphorylation revealed in human cell lines. Nat Chem Biol.

53. Chanduri M., Rai A., Malla A.B., Wu M., Fiedler D., Mallik R., et al. (2016) Inositol hexakisphosphate kinase 1 (IP6K1) activity is required for cytoplasmic dynein-driven transport. Biochem J 473, 3031–47.

54. Jadav R.S., Kumar D., Buwa N., Ganguli S., Thampatty S.R., Balasubramanian N., et al. (2016) Deletion of inositol hexakisphosphate kinase 1 (IP6K1) reduces cell migration and invasion, conferring protection from aerodigestive tract carcinoma in mice. Cell Signal 28, 1124–36.

55. Bhandari R., Juluri K.R., Resnick A.C. and Snyder S.H. (2008) Gene deletion of inositol hexakisphosphate kinase 1 reveals inositol pyrophosphate regulation of insulin secretion, growth, and spermiogenesis. Proc Natl Acad Sci U S A 105, 2349–53.

56. Singh J., Ripp A., Haas T.M., Qiu D., Keller M., Wender P.A., et al. (2019) Synthesis of Modified Nucleoside Oligophosphates Simplified: Fast, Pure, and Protecting Group Free. J Am Chem Soc 141, 15013–7.

57. Suksrichavalit T., Yoshimatsu K., Prachayasittikul V., Bulow L. and Ye L. (2010) “Clickable” affinity ligands for effective separation of glycoproteins. J Chromatogr A 1217, 3635–41.

58. Christ J.J. and Blank L.M. (2018) Enzymatic quantification and length determination of polyphosphate down to a chain length of two. Anal Biochem 548, 82–90.

59. Sivandzade F., Bhalerao A. and Cucullo L. (2019) Analysis of the Mitochondrial Membrane Potential Using the Cationic JC-1 Dye as a Sensitive Fluorescent Probe. Bio Protoc 9.

60. Desfougères Y., Wilson M.S.C., Laha D., Miller G.J. and Saiardi A. (2019) ITPK1 mediates the lipid-independent synthesis of inositol phosphates controlled by metabolism. Proc Natl Acad Sci U S A 116, 24551–61.

61. Plitzko B. and Loesgen S. (2018) Measurement of Oxygen Consumption Rate (OCR) and Extracellular Acidification Rate (ECAR) in Culture Cells for Assessment of the Energy Metabolism. Bio Protoc 8, e2850.

